# Extracellular neuroligin–ICAM5 coupling drives dendritic growth via actin remodeling

**DOI:** 10.64898/2026.03.06.709873

**Authors:** Cydric Geyskens, Brigitt Raux, Nuno Apóstolo, Ellis Boonen, Jeroen Vandensteen, Beatriz Marques, João F. Machado, llkem Kumru, Julie Nys, Eline Creemers, Joris Vandenbempt, Keimpe Wierda, Wim Annaert, Jeffrey N. Savas, Joris de Wit, Jonathan Elegheert, Luís F. Ribeiro

## Abstract

Neuroligins (NLGNs) organize neuronal connectivity by engaging a diverse set of interaction partners, yet how extracellular recognition couples to intracellular growth programs remains unclear. Using affinity proteomics, we identify intercellular adhesion molecule-5 (ICAM5), a cell-surface protein localized to dendritic filopodia, as a novel neuroligin interactor. Surface plasmon resonance and cell-based assays demonstrate direct binding between the ICAM5 and NLGN3 extracellular domains and reveal that ICAM5 engages all neuroligin isoforms. ICAM5 is required for NLGN-induced dendritic outgrowth, but the NLGN3–ICAM5 complex does not contribute to synaptogenesis. Mechanistically, ICAM5 sustains PAK–Cofilin signaling and F-actin organization in growth cones, positioning it as a downstream effector that links neuroligin engagement to actin remodeling. Together, these findings define a neuroligin–ICAM5 axis that couples extracellular recognition to intracellular actin remodeling to control neuronal structural development.

## INTRODUCTION

Synapse formation and development in the brain rely on a complex network of cell adhesion molecules (CAMs) that mediate trans-synaptic interactions and organize pre- and postsynaptic specializations. These adhesion systems not only establish synapse identity but also orchestrate intracellular signaling cascades essential for neuronal connectivity and function ^1–5^. Among them, the neuroligin–neurexin axis represents a prototypical adhesion system for synaptic specification. Postsynaptic neuroligins (NLGNs) engage presynaptic neurexins (NRXNs) through interactions modulated by alternative splicing, enabling isoform-specific regulation of excitatory and inhibitory synapses ^6–8^. In addition to trans-synaptic neurexin binding, neuroligin signaling is refined by *cis*-acting modulators. Members of the MAM domain-containing glycosylphosphatidylinositol anchor proteins (MDGAs) bind neuroligins on the postsynaptic membrane and inhibit their interaction with NRXNs, thereby suppressing synaptogenic activity ^9^. MDGA binding overlaps with the neurexin interface on the neuroligin ectodomain (ECD), establishing a competitive mechanism that fine-tunes synapse formation and maintains synaptic balance ^10^. Through this inhibitory regulation, MDGAs act as negative modulators of neuroligin-dependent synaptic specification ^11^.

Neuroligin-3 (NLGN3) is a particularly intriguing member of the neuroligin family because of its dual localization to both excitatory and inhibitory synapses and its strong link to neurodevelopmental disorders ^12–14^. Mutations in *NLGN3*, most notably leading to the R451C substitution, were first identified in families with X-linked autism spectrum disorder (ASD), establishing neuroligins as critical genetic risk factors ^15^. Mouse models carrying the NLGN3 R451C mutation reproduce key behavioral phenotypes, including impaired social communication and increased repetitive behaviors ^16^, and display enhanced inhibitory transmission ^17^ and altered endocannabinoid signaling ^18^. In contrast to its effects in cortex, where inhibition is preferentially enhanced, the R451C mutation strengthens excitatory transmission in hippocampal circuits, increases AMPA receptor-mediated responses, and augments long-term potentiation (LTP), underscoring the region- and circuit-specific impact of NLGN3 on synaptic strength and plasticity ^19^.

Further underscoring the role of NLGN3 as a key regulator of synaptic function, receptor protein tyrosine phosphatase δ (RPTPδ/PTPRD) has emerged as a ‘non-canonical’ presynaptic ligand for NLGN3 alongside its canonical presynaptic NRXN ligands ^20^. RPTPδ splice variants bind NLGN3 at sites overlapping the neurexin interface, forming mutually exclusive adhesion complexes ^20^. *In vivo* disruption of this RPTPδ–NLGN3 pathway alters social behavior, motor learning, and synaptic protein expression, phenocopying behavioral and molecular aspects of the NLGN3 R451C phenotype ^20^. The canonical NLGN3–NRXN and non-canonical NLGN3–RPTPδ pathways make distinct and partially opposing contributions to social motivation and memory formation ^21^.

In a distinct functional and pathogenic context, soluble NLGN3 ECD acts as a mitogen in the tumor microenvironment (TME). Neuronal activity induces proteolytic shedding of NLGN3, which promotes the proliferation of high-grade glioma (HGG) through activation of the PI3K–mTOR pathway and oncogenic gene programs ^22–24^. This mitogenic activity has been linked to engagement of the chondroitin sulfate proteoglycan CSPG4 (NG2) as a receptor, thereby coupling activity-dependent NLGN3 shedding to CSPG4-dependent signaling, ADAM10 activation, and tumor proliferation ^25^.

NLGN3’s diverse synaptic and non-synaptic functions, together with the emergence of non-canonical ligands, suggest that additional extracellular partners may contribute to its functional repertoire. To explore this possibility, we mapped the NLGN3 interactome by performing affinity proteomic screening with the NLGN3 ectodomain in cortical synaptosomes, which identified ICAM5 (also known as telencephalin; TLN) as a new NLGN3 binding partner. ICAM5 is an immunoglobulin superfamily (IgSF) adhesion molecule enriched in cortical and hippocampal neurons, where it localizes to dendritic filopodia and regulates dendritic growth and spine maturation ^26^. We demonstrate that ICAM5 directly binds all neuroligin isoforms and that ICAM5 is required for neuroligin-driven dendritic outgrowth. Our findings define a neuroligin–ICAM5 signaling axis that links extracellular adhesion to PAK–Cofilin–dependent actin remodeling during neuronal structural development.

## RESULTS

### ICAM5 is a binding partner of NLGN3

To identify previously unrecognized extracellular binding partners of NLGN3, we performed affinity proteomic screening using purified recombinant Fc-linked NLGN3 ectodomain (NLGN3_ECD_-Fc) as bait to isolate interacting proteins from detergent-solubilized synaptosomes prepared from whole rat brain, followed by mass spectrometry analysis ^27^. Among the curated set of cell-surface proteins recovered in the NLGN3 pull-down, intercellular adhesion molecule 5 (ICAM5), a dendritically enriched adhesion molecule, emerged as one of the most prominent candidates, represented by 21 spectral counts (Figure 1A and Table S1). To independently validate this interaction, we performed reciprocal pull-down assays using recombinantly purified ICAM5_ECD_-Fc as bait. In this interactome, NLGN3 was detected with 7 spectral counts, consistent with a NLGN3–ICAM5 protein–protein interaction (Figure 1A and Table S1). The known NLGN3-interacting protein RPTPδ was identified in both pull-downs, providing internal validation of the experimental strategy. In contrast, NRXNs were detected in the NLGN3, but not in the ICAM5, dataset and therefore do not feature in the intersection analysis. MDGA family members were not detected in the NLGN3 dataset.

**Figure 1.**
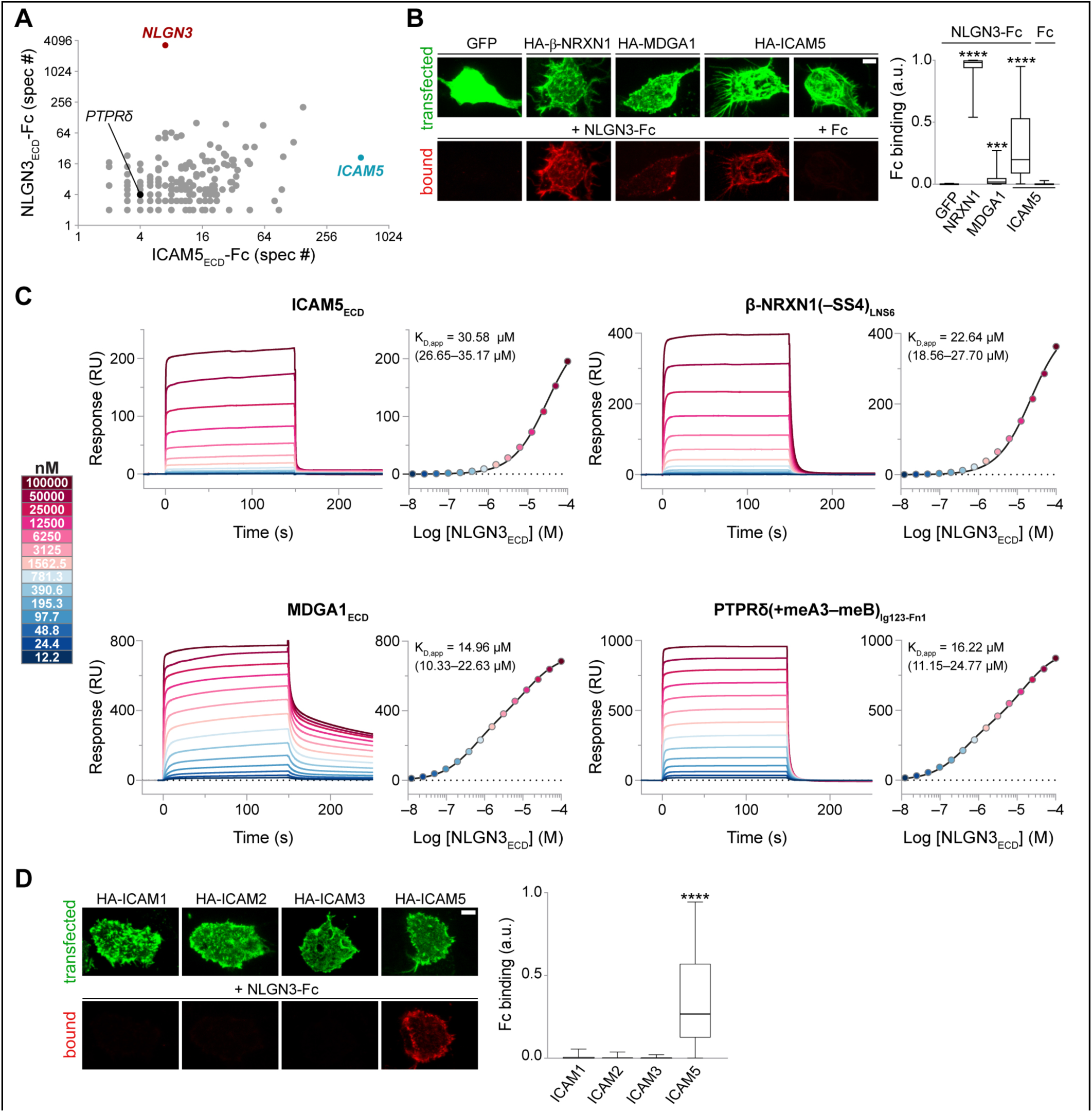
Identification of ICAM5 as a NLGN3-interacting protein. (**A**) Frequency of peptide detection (total spectral counts) for proteins identified in NLGN3ECD-Fc and ICAM5ECD-Fc affinity purifications from detergent-solubilized synaptosomes prepared from post-natal day 21–23 rat brain, following subtraction of proteins detected in Fc-only controls. ICAM5 was detected with 21 spectral counts in the NLGN3-Fc dataset and 548 spectral counts in the ICAM5-Fc dataset, whereas NLGN3 was identified with 3,308 spectral counts in the NLGN3-Fc dataset and 7 spectral counts in the ICAM5-Fc dataset. Each dot represents a protein detected in both datasets and is plotted according to its spectral counts in the respective pull-downs. (**B**) Cell-surface binding assay. NLGN3ECD-Fc (red) bound to HEK293T cells (green) expressing HA-tagged ICAM5, as well as to cells expressing HA-tagged β-NRXN1 or MDGA1 (positive controls). By contrast, NLGN3ECD-Fc did not bind to GFP-expressing cells, and Fc did not bind to ICAM5-expressing cells (negative controls). Scale bar, 5 µm. For quantification, Fc clustering area was normalized to the area of the respective HEK293T cell, as determined by GFP or HA staining (Fc binding). Data are shown as box plots (median, 25^th^–75^th^ interquartile range, and minimum to maximum). ***p = 0.0002; ****p < 0.0001 (Kruskal–Wallis test followed by Dunn’s multiple comparisons test, 2-6 independent experiments). a.u., arbitrary units. (**C**) Surface plasmon resonance (SPR) analysis of direct binding between human NLGN3 ectodomain (NLGN3ECD) and its interaction partners. Recombinant dimeric NLGN3ECD was injected as a two-fold dilution series (100 μM–12 nM) over surface-immobilized human β-NRXN1(–SS4)LNS6, MDGA1ECD, PTPRδ(+meA3–meB)Ig123–Fn1, or ICAM5ECD. Representative sensorgrams (left) and corresponding steady-state binding isotherms (right) are shown. Apparent dissociation constants (KD,app, mean with 95% confidence intervals) are annotated. (**D**) Cell-surface binding assay. NLGN3ECD-Fc (red) bound to HEK293T cells (green) expressing HA-tagged ICAM5, but not to cells expressing the ICAM1, ICAM2, or ICAM3 ECDs. Scale bar, 5 µm. For quantification, Fc clustering area was normalized to the area of the respective HEK293T cell, as determined by HA staining (Fc binding). Data are shown as box plots (median, 25th–75th percentile, and minimum to maximum). ****p < 0.0001 (Kruskal–Wallis test followed by Dunn’s multiple comparisons test, 3 independent experiments). a.u., arbitrary units. See also Figure S1.

To corroborate the proteomic results and investigate a direct NLGN3–ICAM5 interaction, we performed cell-surface binding and surface plasmon resonance (SPR) assays. First, we expressed the membrane-anchored, hemagglutinin (HA)-tagged full-length (Ig1–9) ICAM5 ECD in HEK293T cells. We then applied NLGN3_ECD_-Fc or Fc-only (as a negative control) and assessed Fc binding to the cell surface by immunocytochemistry. In control experiments, HEK293T cells were transfected with plasmids encoding the NLGN3-binding partners β-NRXN1 or MDGA1, or with green fluorescent protein (GFP), and then treated with NLGN3_ECD_-Fc. NLGN3_ECD_-Fc bound to β-NRXN1-, MDGA1- and ICAM5-expressing cells, whereas no detectable binding was observed to GFP-expressing cells or to ICAM5-expressing cells treated with Fc (Figure 1B), supporting a direct interaction between the NLGN3 and ICAM5 ECDs.

We next performed quantitative SPR analyses using purified recombinant ECDs to determine equilibrium binding affinities. Concentration series of dimeric NLGN3_ECD_ (100 μM–12.2 nM) were injected over surface-immobilized full-length ICAM5 ECD, β-NRXN1 LNS6 domain (lacking the splice site 4 (SS4) insert; –SS4), PTPRδ_Ig123-Fn1_, or MDGA1 ECD (Figure 1C). In line with previous results ^10^, NLGN3 bound β-NRXN1(–SS4) with an apparent dissociation constant (K_D,app_) of ∼22 μM, while it interacted with MDGA1 and PTPRδ with comparable apparent affinities of ∼15–16 μM. Direct binding to ICAM5 was clearly detectable, yet also notably weak with a K_D,app_ of ∼30 μM (95 % CI: 26.6–35.2 μM) and concomitant very fast dissociation kinetics. These results establish ICAM5 as a direct, yet low-affinity binding partner of NLGN3, and reveal a consistent pattern in which all currently known extracellular binding partners of NLGN3 interact transiently, a feature that may enable dynamic partner exchange and context-dependent signaling at neuronal membranes.

We then investigated which of the nine Ig domains of ICAM5 (Ig1–9) mediate the NLGN3–ICAM5 interaction. In HEK293T cells expressing ICAM5 Ig1 or Ig1–3, we observed intermediate binding levels compared with full-length ICAM5 (Ig1–9), whereas no binding was detected in cells expressing Ig2, Ig3, Ig2–3, or Ig4–9 (Figures S1A–B). These results indicate that Ig1 is required for the NLGN3–ICAM5 interaction but is not sufficient to fully recapitulate the binding observed with the full-length ECD, suggesting that its intact architecture is required for optimal interaction. Notably, integrin αLβ2 engages the N-terminal Ig1–2 domains of ICAM5 ^28^, indicating that the membrane-distal portion of ICAM5 serves as a shared recognition platform for its distinct extracellular partners.

Finally, to assess whether NLGN3 may interact with the other neuronal ICAMs, we performed cell-surface binding assays by expressing the membrane-anchored ECDs of ICAM1, ICAM2, ICAM3, or ICAM5 (Figure 1D). ICAM4 was not included because it is an erythroid-restricted adhesion molecule and is not expressed in the nervous system. Binding was detected only with ICAM5 (Figures 1D), indicating that NLGN3 selectively recognizes ICAM5.

### All neuroligins can bind to ICAM5

To determine whether ICAM5 recognition is unique to NLGN3 or shared across the neuroligin family, we performed SPR experiments using the ECDs of NLGN1–3, NLGN4X, and NLGN4Y. In contrast to the initial configuration in which immobilized ICAM5 was probed with dimeric NLGN3 under avidity-permissive conditions (Figure 1C), these experiments employed immobilized dimeric neuroligins and monomeric analytes, thereby minimizing avidity contributions and more closely reflecting intrinsic binding affinities (Figure 2). Each neuroligin was probed with serial dilutions (50 μM–12.2 nM) of recombinant ICAM5_ECD_ (Ig1–9), β-NRXN1(–SS4)_LNS6_ or MDGA1_ECD_, which yielded clear concentration-dependent binding responses across all neuroligin isoforms. ICAM5 displayed notably weaker interactions than either β-NRXN1 or MDGA1 with all NLGNs, with K_D_s > 10 μM (Figure 2). Interestingly, NLGN1 and NLGN2 bound ICAM5 more robustly than NLGN3, based on apparent binding responses. Among the neuroligins, NLGN1 and NLGN2 also interact strongest with β-NRXN1 (K_D_ ∼0.8 µM) and MDGA1 (K_D_ ∼1.3 µM), establishing a consistent affinity pattern across these ligands. Despite their high abundance in brain tissue, NLGN1 and NLGN2 were however not recovered in the reciprocal ICAM5 pull-downs (Figure 1A and Table S1), indicating that ICAM5 association with these isoforms may be limited under native conditions. In summary, although ICAM5 engages all neuroligins, the interaction is substantially weaker than canonical NRXN or MDGA binding and varies across neuroligin isoforms.

**Figure 2.**
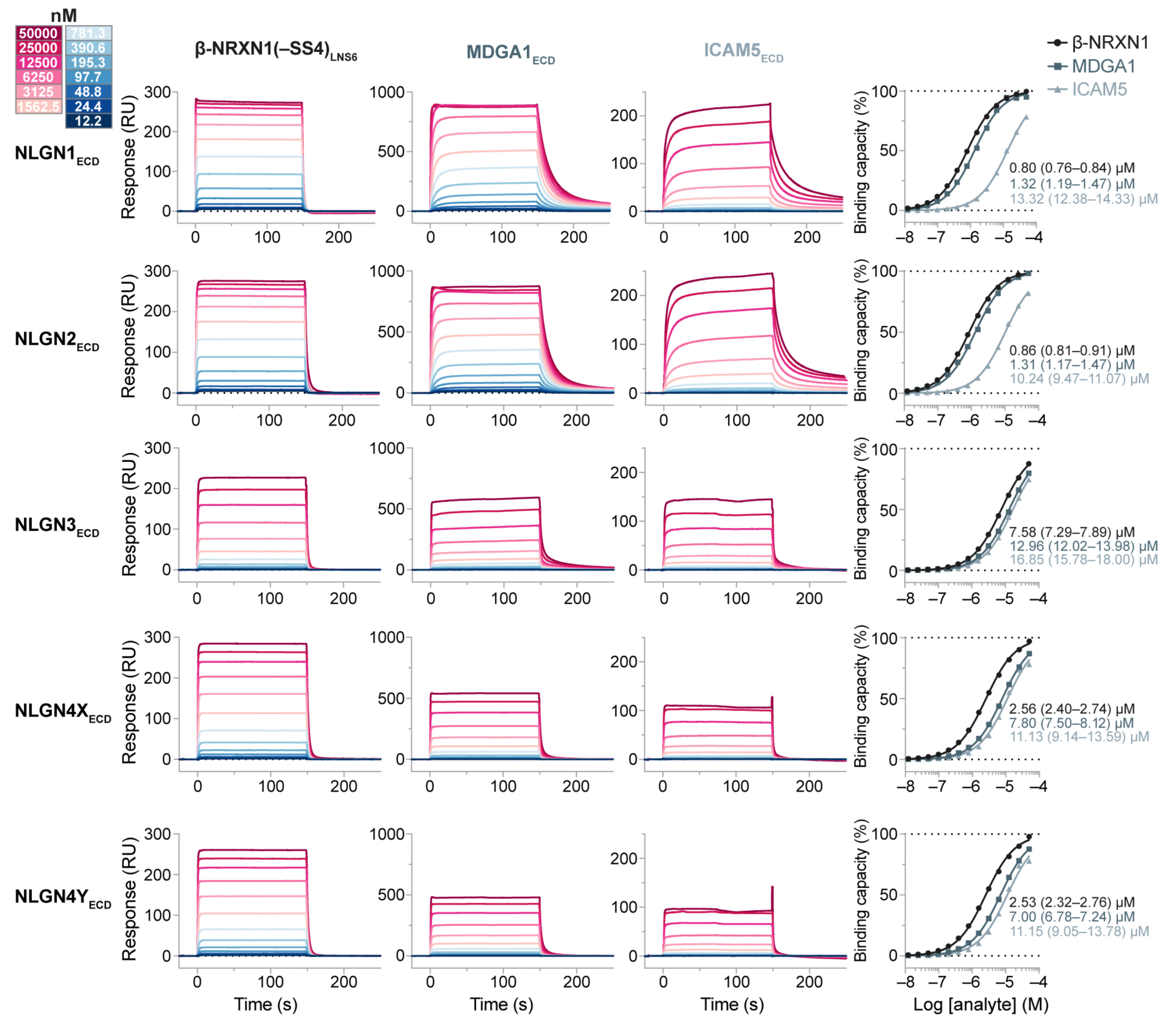
All neuroligin ectodomains can bind the ICAM5 ectodomai. Ectodomains of the human neuroligins (NLGN1, NLGN2, NLGN3, NLGN4X, and NLGN4Y) were surface-immobilized probed with two-fold dilution series (50 μM–12 nM) of human β-NRXN1(–SS4)LNS6, MDGA1ECD, or ICAM5ECD, yielding representative SPR sensorgrams (left) and corresponding normalized binding isotherms (right) for each neuroligin isoform. Binding affinities (KD, mean with 95% confidence intervals) are annotated.

### NLGN3–ICAM5 is not involved in the control of synapse formation

Because both NLGN3 and ICAM5 have been implicated in excitatory synapse development, NLGN3 as a positive regulator ^29,30^ and ICAM5 as a negative modulator ^31^ of synapse maturation, we first asked whether the NLGN3–ICAM5 interaction contributes to the regulation of excitatory synapse density. To address this, we expressed GFP together with either an empty vector (control), NLGN3, or ICAM5 in cortical neurons prepared from wild-type (WT) or *Icam5* knockout (KO) littermates, as well as from WT or *Nlgn3* KO littermates, and quantified the fluorescence intensity of the excitatory postsynaptic marker PSD-95. Consistent with previous reports, overexpression of NLGN3 increased PSD-95 intensity ^29,30^, whereas ICAM5 overexpression reduced it. Conversely, genetic deletion of either molecule produced trends opposite to their respective overexpression phenotypes (Figures S2A–D). Importantly, these effects were not altered by deletion of the other protein, indicating that NLGN3 and ICAM5 regulate excitatory synapse density independently.

To further examine whether ICAM5 modulates the synaptogenic activity of neuroligins, we compared its effect with that of MDGA1, which inhibits neuroligin-induced presynaptic differentiation by blocking interactions with neurexins ^9,10^. We therefore performed a heterologous synaptogenic co-culture assay in which HEK293T cells expressing NLGN3 or NLGN1 together with either an empty vector (control), ICAM5, or MDGA1 were co-cultured with cortical neurons, and recruitment of Synapsin-positive presynaptic terminals onto HEK293T cells was quantified. As expected ^10^, co-expression of MDGA1 significantly reduced the synaptogenic activity of neuroligins. In contrast, ICAM5 co-expression had no detectable effect on the ability of NLGN1 or NLGN3 to recruit Synapsin-positive presynaptic terminals (Figures S2E–G). Together, these findings indicate that ICAM5 does not modulate neuroligin-induced presynaptic differentiation and therefore does not appear to participate in the canonical synaptogenic pathway mediated by neuroligin–neurexin interactions.

### Neuroligins promote dendritic outgrowth through ICAM5

Having ruled out the involvement of the NLGN3–ICAM5 interaction in excitatory synapse formation, we next asked whether this interaction might instead influence earlier stages of neuronal development. Because both NLGN3 and ICAM5 have previously been implicated in dendritic morphogenesis ^32–35^, we examined whether the NLGN3–ICAM5 complex contributes to dendritic outgrowth. To this end, we overexpressed GFP together with either an empty vector (control), NLGN3, or ICAM5 in cortical neurons prepared from WT or *Icam5* KO littermates, as well as from WT or *Nlgn3* KO littermates, and analyzed total dendritic length using GFP signal in microtubule-associated protein 2 (MAP2)-positive neurites. Overexpression of NLGN3 increased the total dendritic length in WT neurons. In *Icam5* KO neurons, NLGN3 overexpression still promoted dendritic growth but to a far lesser extent than in WT neurons, indicating that ICAM5 enhances NLGN3-induced dendritic outgrowth (Figures 3A–B). We next asked whether NLGN3 is required for ICAM5-mediated dendritic growth. Overexpression of ICAM5 robustly increased dendritic length in both WT and Nlgn3 KO neurons (Figures 3C–D), indicating that ICAM5 can promote dendritic outgrowth independently of NLGN3.

**Figure 3.**
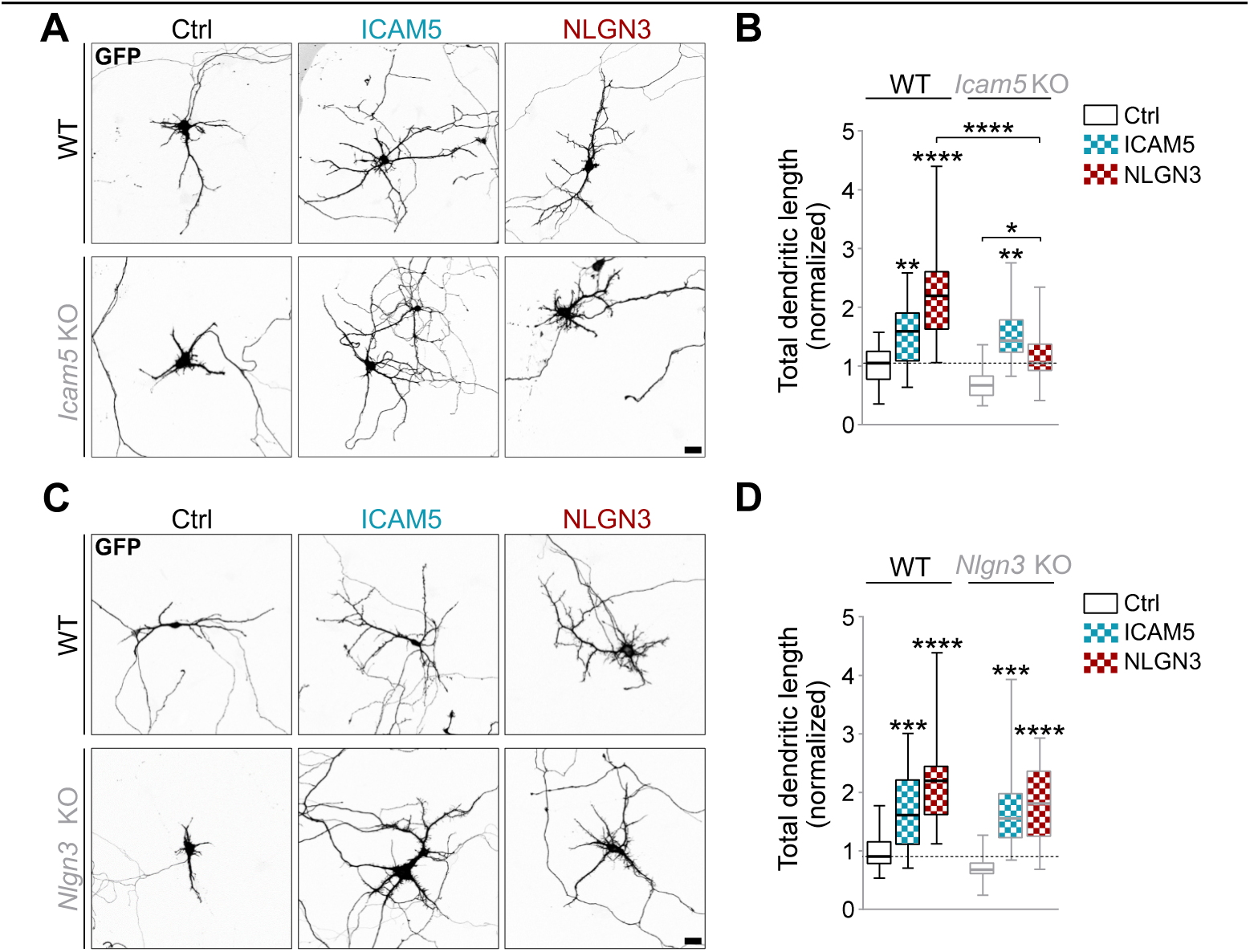
ICAM5 is required for NLGN3-induced dendritic growth. (**A**) Representative images of DIV 7 WT or *Icam5* KO mouse cortical neurons transfected at DIV 3 with GFP and empty vector (Ctrl), GFP and FLAG-tagged ICAM5, or GFP and HA-tagged NLGN3. Neurons were immunostained for GFP (grayscale), MAP2 (not shown), and HA or FLAG (not shown). Scale bar, 20 µm. (**B**) Quantification of panel A: total dendritic length, defined as GFP signal in MAP2-positive neurites and normalized to control cells. Data are shown as box plots (median, 25^th^–75^th^ percentile, and minimum to maximum). *p = 0.0142; **p < 0.01; ****p < 0.0001 (Kruskal–Wallis test followed by Dunn’s multiple comparisons test, 3 independent experiments). (**C**) Representative images of DIV 7 WT or *Nlgn3* KO mouse cortical neurons transfected at DIV 3 with GFP and empty vector (Ctrl), GFP and FLAG-tagged ICAM5, or GFP and HA-tagged NLGN3. Neurons were immunostained for GFP (grayscale), MAP2 (not shown), and HA or FLAG (not shown). Scale bar, 20 µm. (**D**) Quantification of panel C: total dendritic length, defined as GFP signal in MAP2-positive neurites and normalized to control cells. Data are shown as box plots (median, 25^th^–75^th^ percentile, and minimum to maximum). ***p < 0.001; ****p < 0.0001; (Kruskal–Wallis test followed by Dunn’s multiple comparisons test, 3 independent experiments). See also Figure S3.

Because we found that multiple neuroligin isoforms interact with ICAM5, we next asked whether additional neuroligins also promote dendritic outgrowth in an ICAM5-dependent manner. We focused on NLGN4, which has previously been implicated in dendritic growth regulation ^34^. We therefore overexpressed GFP together with either an empty vector (control), NLGN4X, NLGN4Y, or ICAM5 in cortical neurons prepared from WT or *Icam5* KO littermates and quantified total dendritic length as described above. Similar to NLGN3, overexpression of NLGN4X increased dendritic length in WT neurons but produced a markedly reduced effect in *Icam5* KO neurons (Figures S3A–B), indicating that ICAM5 also contributes to NLGN4X-induced dendritic outgrowth. In contrast, NLGN4Y overexpression did not significantly increase dendritic growth (Figures S3A–B). In our experimental conditions, NLGN4Y indeed exhibited lower surface expression compared with NLGN3 and NLGN4X (Figure S3C), consistent with previous reports ^36^.

Together, these results indicate that neuroligins promote dendritic outgrowth through mechanisms that are strongly enhanced by ICAM5 and place ICAM5 downstream of neuroligin signaling in the regulation of dendritic growth.

### ICAM5 links neuroligins to PAK–Cofilin signaling and actin cytoskeleton organization

To investigate the mechanisms by which the NLGN3–ICAM5 complex regulates dendritic growth, we examined the effects of NLGN3 or ICAM5 loss-of-function on intracellular signaling pathways linked to neuroligin-dependent control of neuronal growth and synaptic plasticity, including protein translation ^37^, mTOR signaling, PI3K–Akt signaling, and PAK-dependent actin regulation ^22,34,35^. We first assessed global protein synthesis using the SUnSET assay, in which incorporation of puromycin into nascent polypeptides provides a quantitative readout of translational activity. In parallel, we measured phosphorylation levels of 4E-BP1, Akt, and PAK as readouts of mTOR-, PI3K-, and PAK-dependent signaling, respectively (Figures 4A–D and S4A–D). Both *Nlgn3* KO and *Icam5* KO cortical neurons exhibited reduced global protein synthesis compared with WT neurons (Figure S4A–B). Consistent with this observation, phosphorylation of 4E-BP1 (Thr37/46) was decreased in both mutant conditions, indicating reduced mTOR-dependent translational activity. Analysis of upstream signaling revealed distinct pathway perturbations in the two knockout models. In *Nlgn3* KO neurons, phosphorylation of PAK1/2 (Ser144/141) was reduced, whereas phosphorylation of Akt (Ser473) was increased (Figures S4C–D), suggesting compensatory activation of PI3K–Akt signaling. In contrast, *Icam5* KO neurons displayed reduced phospho-4E-BP1 levels together with a marked reduction in total PAK protein levels (Figures 4A–B), indicating impaired mTOR signaling and destabilization of PAK.

**Figure 4.**
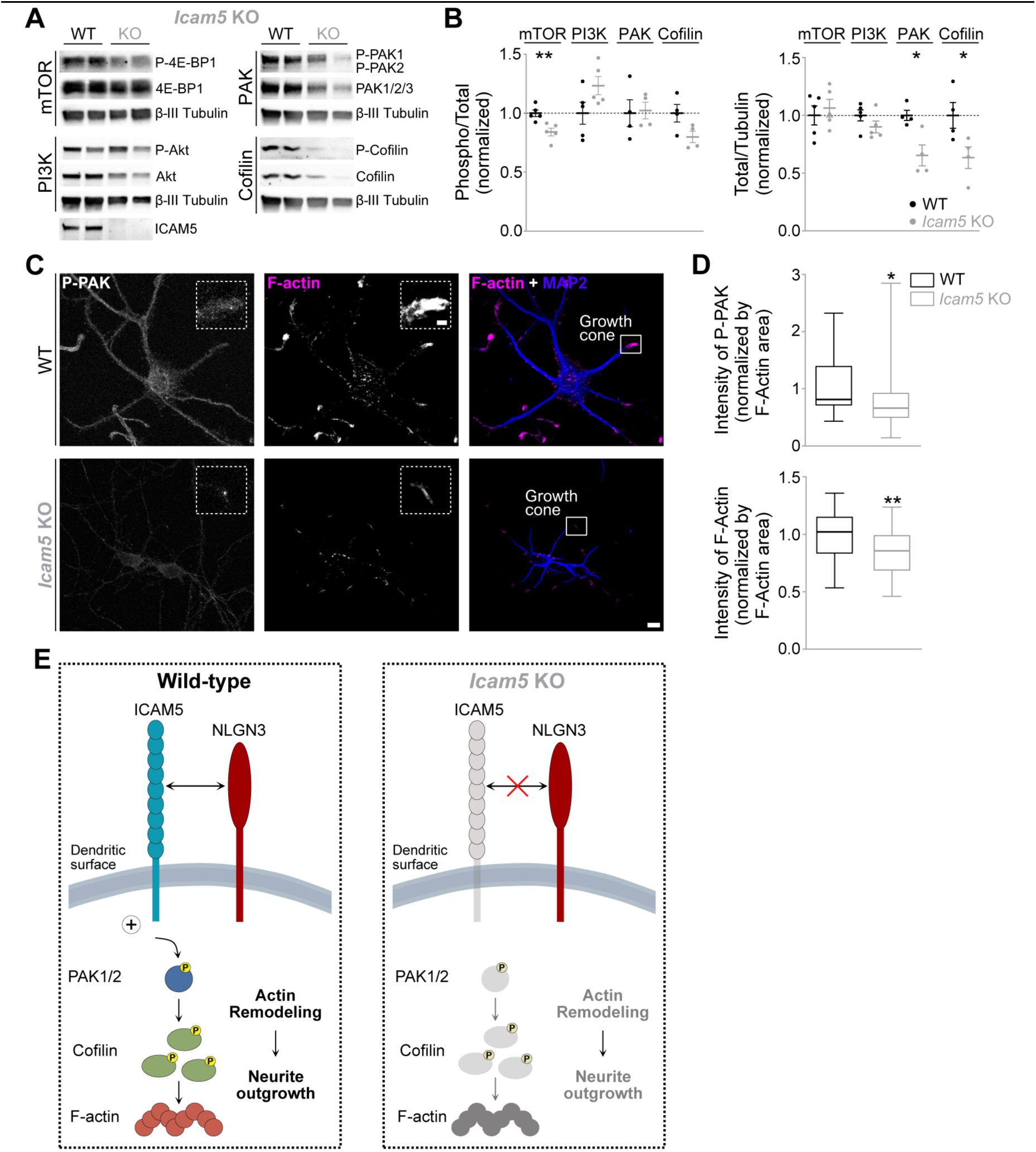
ICAM5 is required for PAK–Cofilin signaling and F-actin stabilization. (**A**) Western blot analysis of signaling pathways in DIV7 cortical neurons derived from WT or *Icam5* KO mice. Lysates were probed for markers of mTOR signaling (4E-BP1 and phospho-4E-BP1 [Thr37/46]), PI3K signaling (Akt and phospho-Akt [Ser473]), PAK signaling (total PAK1–3 and phospho-PAK1 [Ser144] / phospho-PAK2 [Ser141]), and the actin regulator Cofilin (total and phospho-Cofilin [Ser3]). βIII-tubulin was used as a loading control, and ICAM5 immunoblotting confirmed loss of ICAM5 expression in *Icam5* KO neurons. (**C**) Quantification of panel A: phospho-band intensities were normalized to the corresponding total protein levels and to WT control, while total protein band intensities were normalized to βIII-tubulin and to WT control. Data are shown as individual data points with mean ± SEM. *p < 0.05; **p < 0.01 (Unpaired t-test, 4-5 independent experiments).Representative images of DIV7 WT or *Icam5* KO mouse cortical neurons immunostained for phospho-PAK (p-PAK1 Ser144 / p-PAK2 Ser141; grayscale), F-actin (phalloidin; magenta and grayscale), and MAP2 (blue). Insets show magnified views of selected growth cones. Scale bars, 10 µm and 2 µm (insets). (**D**) Quantification of panel C: Quantification of phospho-PAK or F-actin intensity within dendritic growth cones, defined by phalloidin staining (F-actin) in MAP2-positive neurites, normalized to F-actin area and to WT control. Data are shown as box plots (median, 25^th^–75^th^ percentile, and minimum to maximum). *p = 0.0426 (Mann–Whitney test, 3 independent experiments; phospho-PAK) and *p = 0.0018 (Unpaired t-test, 3 independent experiments; F-actin). (**E**) Schematic model illustrating how NLGN3 and ICAM5 cooperate at the dendritic surface to regulate intracellular signaling and actin cytoskeleton organization during neuronal development. In wild-type neurons, interaction between NLGN3 and ICAM5 promotes PAK1/2 signaling and downstream phosphorylation of Cofilin, supporting F-actin assembly and dendritic outgrowth. In Icam5 knockout (Icam5 KO) neurons, loss of ICAM5 impairs neuroligin-driven signaling, resulting in reduced PAK1/2 and Cofilin phosphorylation, decreased F-actin levels, and diminished neurite outgrowth. See also Figure S4.

Because both ICAM5 and PAK have been linked to regulation of the actin-binding protein Cofilin ^38,39^, we next examined total and phosphorylated Cofilin levels. Total Cofilin levels were reduced in *Icam5* KO neurons but remained unchanged in *Nlgn3* KO neurons (Figures 4A–B and S4C–D), supporting a role for ICAM5 in maintaining the stability of both PAK and Cofilin. Consistent with these biochemical changes, phalloidin staining revealed reduced phospho-PAK and F-actin signals within dendritic growth cones of *Icam5* KO neurons (Figures 4C–D). Together, these findings indicate that ICAM5 links growth-promoting neuroligin signaling to the PAK–Cofilin pathway that regulates actin cytoskeleton organization during dendritic growth. In agreement with this model, loss of ICAM5, but not NLGN3, destabilized PAK and Cofilin and impaired neuroligin-induced dendritic outgrowth.

## DISCUSSION

We identify ICAM5 (telencephalin) as a direct, low-affinity binding partner for NLGN3 and, more broadly, for all human neuroligins. Functionally, the NLGN–ICAM5 axis promotes dendritic outgrowth without impacting synaptogenesis, separating growth control from classical synapse-organizing roles. This expands the extracellular recognition repertoire of neuroligins beyond NRXNs, MDGAs and RPTPδ. ICAM5’s dendritic localization, enrichment on filopodia, regulated ectodomain shedding, and links to β1-integrins and cytoskeletal adaptors provide a coherent route by which neuroligin engagement can access actin machinery. Our data connect neuroligin binding to PAK–Cofilin signaling and growth-cone F-actin organization and support a model in which neuroligin signaling is encoded by modular extracellular recognition programs: (i) synaptogenic specification mediated by NRXNs; (ii) non-canonical presynaptic coupling via RPTPδ; (iii) MDGA-dependent restraint of synaptogenesis; and (iv) ICAM5-dependent control of dendritic growth. Notably, the expression patterns of ICAM5 and neuroligins during neuronal development are consistent with such a role. ICAM5 is highly enriched in dendritic filopodia during early stages of dendritic growth and declines as synapses mature ^40,41^, whereas neuroligins are expressed during neuronal maturation ^42^, suggesting that NLGN–ICAM5 interactions may operate during a developmental window that links dendritic morphogenesis to emerging synaptic connectivity.

### ICAM5 promotes dendritic growth by coupling cell-surface adhesion to intracellular signaling

ICAM5 is a strictly somatodendritic adhesion molecule ^40,43^ that is enriched in dendritic filopodia and growth cones ^40,44–46^. This distinctive localization pattern makes ICAM5 particularly relevant for neuronal development, especially during dendritic outgrowth. Indeed, substrates coated with recombinant ICAM5 promote dendrite growth, an effect mediated, at least in part, by homophilic ICAM5 interactions ^33^.

Consistent with these observations, we show here that ICAM5 overexpression increases total dendritic length (Figure 3).

Beyond the contribution of homophilic interactions, ICAM5-mediated dendritic outgrowth depends on signaling mediated by its cytoplasmic tail, which interacts with members of the ezrin–radixin–moesin (ERM) family of cytoskeletal linker proteins ^44^. These proteins associate in turn with actin filaments and function as cross-linkers that connect the plasma membrane to the underlying cytoskeleton ^47,48^. Notably, ERM proteins were also detected in the ICAM5 interactome (Figure S4E and Table S1), supporting their association with ICAM5-containing complexes. ICAM5 colocalizes with both α-actinin and F-actin, and disruption of the ICAM5–α-actinin interaction impairs the ability of ICAM5 to induce dendritic outgrowth ^49^.

Supporting a role for ICAM5 as a molecular linker between extracellular adhesion and the actin cytoskeleton, we show that ICAM5-deficient neurons exhibit reduced F-actin levels within dendritic growth cones. Moreover, these neurons display decreased total levels of Cofilin and PAK (Figure 4), key regulators of actin dynamics ^50,51^. Consistent with this observation, our ICAM5 interactome analysis identified PAK1–3 and Cofilin 1 as ICAM5-associated proteins (Figure S4E and Table S1). Because the interactome was performed using the ICAM5 ECD, these interactions may arise from endogenous ICAM5 forming homophilic interactions with the recombinant ICAM5 bait used in the assay. At present, it remains unclear whether the association of ICAM5 with PAK and Cofilin is direct or mediated through additional adaptor proteins. Nonetheless, these findings support a model in which ICAM5 acts as a signaling hub at the dendritic plasma membrane, coordinating extracellular adhesion cues with intracellular pathways that regulate actin cytoskeleton organization.

Taken together, our data support a model in which ICAM5 promotes dendritic growth by integrating cell-surface adhesion with actin remodeling through multiple, potentially convergent mechanisms. These include direct interactions with actin-binding proteins such as ERM proteins and α-actinin, as well as the modulation of actin-regulatory signaling pathways involving PAK and Cofilin. Through these combined actions, ICAM5 is well positioned to locally control cytoskeletal organization within dendritic growth cones and thereby drive dendritic morphogenesis.

### Neuroligins regulate neurite outgrowth

Similar to ICAM5, neuroligins, particularly NLGN3 and NLGN4X, have been implicated in the regulation of cellular growth processes, including neurite outgrowth ^34,35^. Loss-of-function studies in primary hippocampal cultures have shown that shRNA-mediated knockdown of NLGN3 reduces dendritic outgrowth ^35^. Conversely, overexpression of NLGN3 or NLGN4X promotes neurite outgrowth in primary cortical neurons as well as in undifferentiated human neural progenitor cells ^34^. Consistent with these observations, the results presented here further support a role for neuroligins as positive regulators of dendritic outgrowth.

Accumulating evidence indicates that manipulation of NLGN3 expression levels in neurons modulates multiple intracellular signaling pathways, including PI3K, mTOR, PAK, and Cofilin ^34,35,37^, which are known to influence cellular growth. In agreement with these studies, we detected alterations in PI3K, mTOR, and PAK signaling following NLGN3 loss-of-function (Figures S4C–D). Such changes may influence neurite outgrowth through multiple mechanisms, including regulation of protein synthesis and actin cytoskeleton remodeling.

Consistent with this framework, overexpression of NLGN3 and NLGN4X has been reported to activate PAK signaling and Cofilin in developing neurons, where both proteins are enriched in neurite growth cones and PAK activity is required for the neurite outgrowth-promoting effects of neuroligins ^34^. Notably, we observe that ICAM5 loss-of-function leads to a marked reduction in PAK and Cofilin levels (Figure 4), suggesting that ICAM5 contributes to neuroligin-dependent activation of these signaling pathways. Together, these findings support a model in which neuroligins promote dendritic outgrowth through ICAM5-dependent coupling of extracellular adhesion to intracellular signaling pathways that regulate actin dynamics and neuronal growth (Figure 4E).

More broadly, these observations align with the concept of synaptotropic dendritic growth, in which dendritic branching and stabilization are influenced by synaptic adhesion cues during early neuronal development. In this context, neuroligins may contribute to dendritic morphogenesis not only through their established roles in synapse formation but also by locally regulating cytoskeletal dynamics at developing dendritic growth cones ^52^.

Beyond its role in regulating intracellular signaling during neuronal development, neuron-derived secreted NLGN3 has been shown to promote glioma cell proliferation through activation of the PI3K–mTOR signaling pathway ^22,23^, highlighting the broader capacity of NLGN3 to engage growth-promoting signaling cascades across distinct cellular contexts. Whether ICAM5 is required for the activation of signaling pathways that promote NLGN3-dependent glioma cell proliferation remains an open question.

### The neuroligin ectodomain acts as a hub for diverse extracellular interactions

The NRXN, RPTPδ, and MDGA proteins had previously been identified as neuroligin interactors. Here, we show that neuroligins also directly engage ICAM5, expanding the repertoire of extracellular partners associated with neuroligins. All human neuroligin isoforms bound ICAM5, indicating that this interaction represents a conserved feature of the neuroligin family. Conversely, ICAM5 was uniquely capable of binding neuroligins within the ICAM family, highlighting the specificity of this interaction.

Together, these findings support a model in which neuroligins act as modular extracellular recognition hubs that recruit distinct partners to couple adhesion events to intracellular signaling pathways. By identifying ICAM5 as a neuroligin-binding partner that links neuroligin adhesion to PAK–Cofilin–dependent actin regulation, our work expands the functional scope of neuroligin signaling beyond synapse formation to include the control of dendritic growth during neuronal development. More broadly, these findings illustrate how cell adhesion molecules integrate extracellular recognition with intracellular signaling to shape neuronal architecture.

### Limitations of the study

Our biophysical conclusions derive from SPR with recombinant ectodomains; accordingly, the reported K_D_ values reflect low-affinity binding capacity *in vitro* and may not translate to the in-membrane, *in vivo* context, where two-dimensional confinement, local copy numbers/stoichiometry, membrane organization, and crowding can alter effective association/dissociation rates and avidity. The low affinity of the recombinant NLGN–ICAM5 interaction precluded cryo-EM or crystallographic structure determination, and complex structure predictions did not yield confident or interpretable models. Functional assays relied on overexpression and full knockouts in rodent cortical cultures; we did not perform *in vivo* loss-of-function or knock-in rescue, nor longitudinal analyses of arbor maturation, circuit wiring, or behavior. Mechanistically, the PAK–Cofilin axis was inferred from protein/phospho levels and growth-cone F-actin; causal sufficiency (e.g., PAK rescue) remains to be defined. Finally, we did not perform competition or head-to-head comparisons between NLGNs and other established ICAM5 ligands (e.g., β1 integrins), nor competition of ICAM5 *versus* NRXN or RPTPδ for binding to NLGN; thus, while we establish direct binding capacity and growth-related phenotypes, the relative hierarchy, co-engagement, and pathway partitioning remain unresolved and the subject of future investigations.

## RESOURCE AVAILABILITY

### Lead contact

Further information and requests for resources and reagents should be directed to and will be fulfilled by the lead contact, Luís F. Ribeiro.

### Materials and data availability

All unique reagents generated in this study are listed in the key resources table and available from the lead contact with a completed materials transfer agreement. Any additional information required to reanalyze the data reported in this work is available from the lead contact upon request.

## ACKNOWLEDGMENTS

This work was supported by the European Research Council (ERC) Starting Grant 850820 (SynLink) and by the Région Nouvelle-Aquitaine Grant 3428720 to J.E.; by the European Union’s Horizon 2020 research and innovation programme under Grant Agreement No. 857524, by the Marie Skłodowska-Curie Grant Agreement No. 101031398, by national funds through the Portuguese Science and Technology Foundation (FCT: PTDC/BIA-CEL/2286/2020), and by the FWO postdoctoral grant (12N0319N) to L.F.R; by the FWO Project Grants G0C4518N, G0A8720N, G0A8320N, by the FWO EOS Grant G0H2818N, and by ERANET-NEURON TAO2PATHY Grant G0I3118N to J.d.W. We also acknowledge support from the MICC Imaging Facility of CNC-UC, partially funded by the Portuguese Platform of BioImaging (PPBI; PPBI-POCI-01-0145-FEDER-022122). We would like to thank Dr. Nils Brose (Max Planck Institute for Multidisciplinary Sciences) for providing the *Nlgn3* KO mouse line.

## AUTHOR CONTRIBUTIONS

J.d.W., J.E., and L.F.R. conceived the project and designed the experiments. C.G., B.R., N.A., E.B., J.V., B.M., J.F.M., l.K., J.N., E.C., J.V., K.W., and L.F.R conducted and analyzed experiments. J.N.S. performed mass spectrometric analysis. W.A. contributed with reagents. C.G., B.R., N.A., K.W., J.N.S., J.d.W., J.E., and L.F.R analyzed data. J.d.W., J.E., and L.F.R. wrote the paper, with input from all authors.

## DECLARATION OF INTERESTS

J.d.W. is scientific co-founder and served as scientific advisory board member of Augustine Therapeutics.

## SUPPLEMENTAL INFORMATION

Figures S1–S4.

Table S1. The spreadsheets list the proteins identified by affinity proteomics and mass spectrometry following pull-down assays using recombinant NLGN3_ECD_-Fc or ICAM5_ECD_-Fc as bait. The spreadsheets include: (i) proteins detected in the NLGN3-Fc dataset, (ii) proteins detected in the ICAM5-Fc dataset, and (iii) proteins identified in both datasets (intersection analysis), corresponding to those represented in Figure 1A. Spectral counts are provided for each protein in the respective datasets.

## STAR★METHODS

### KEY RESOURCES TABLE

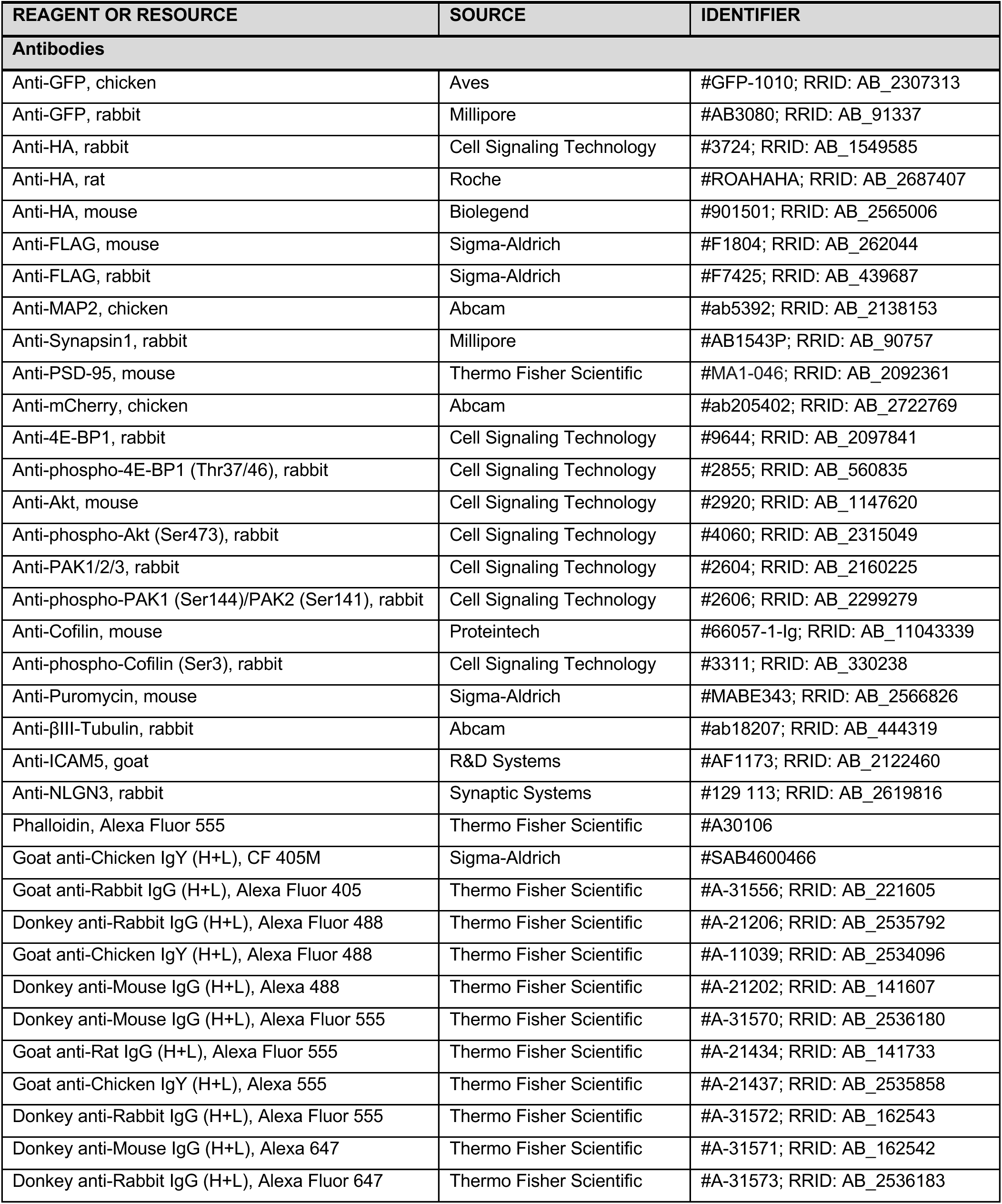
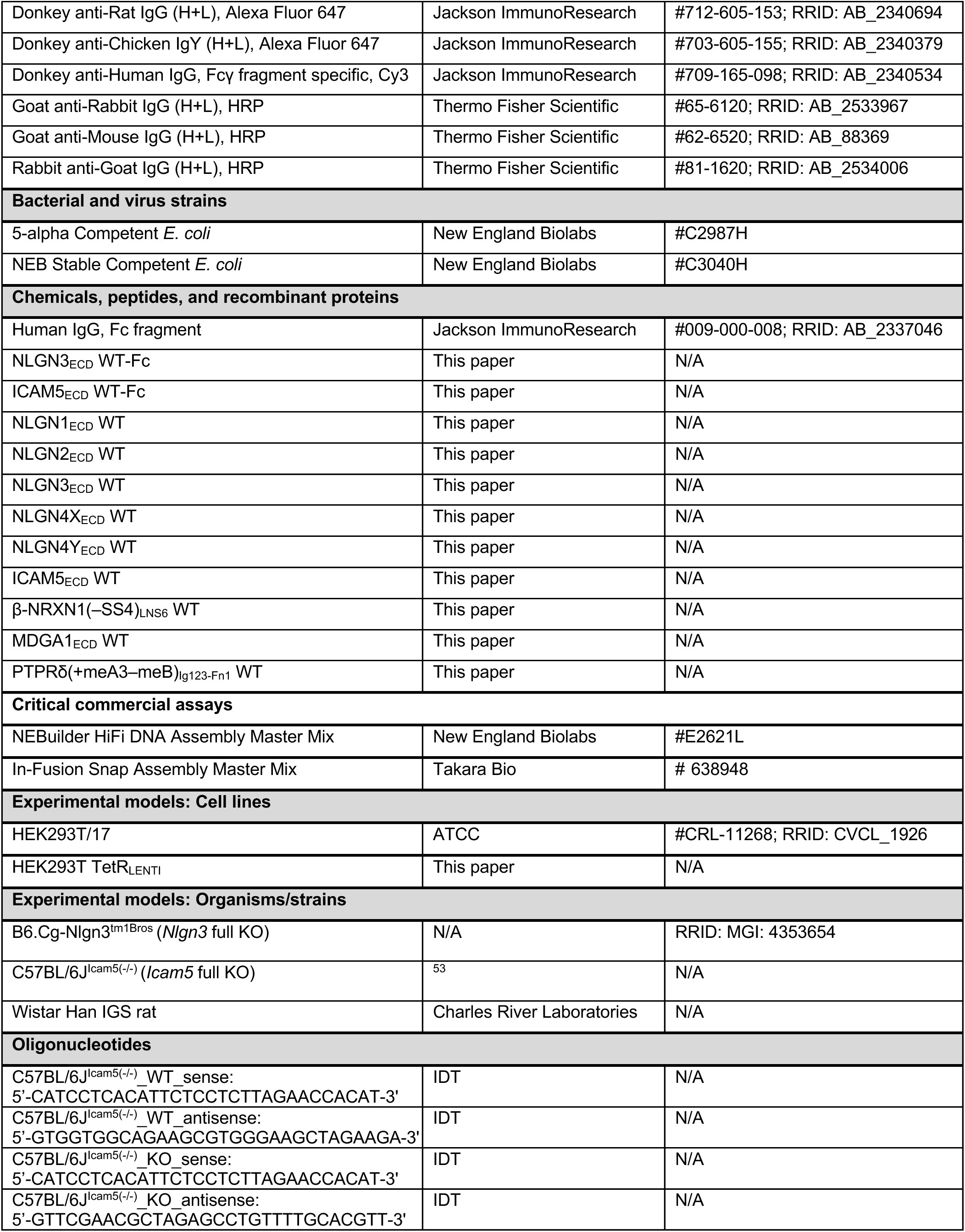
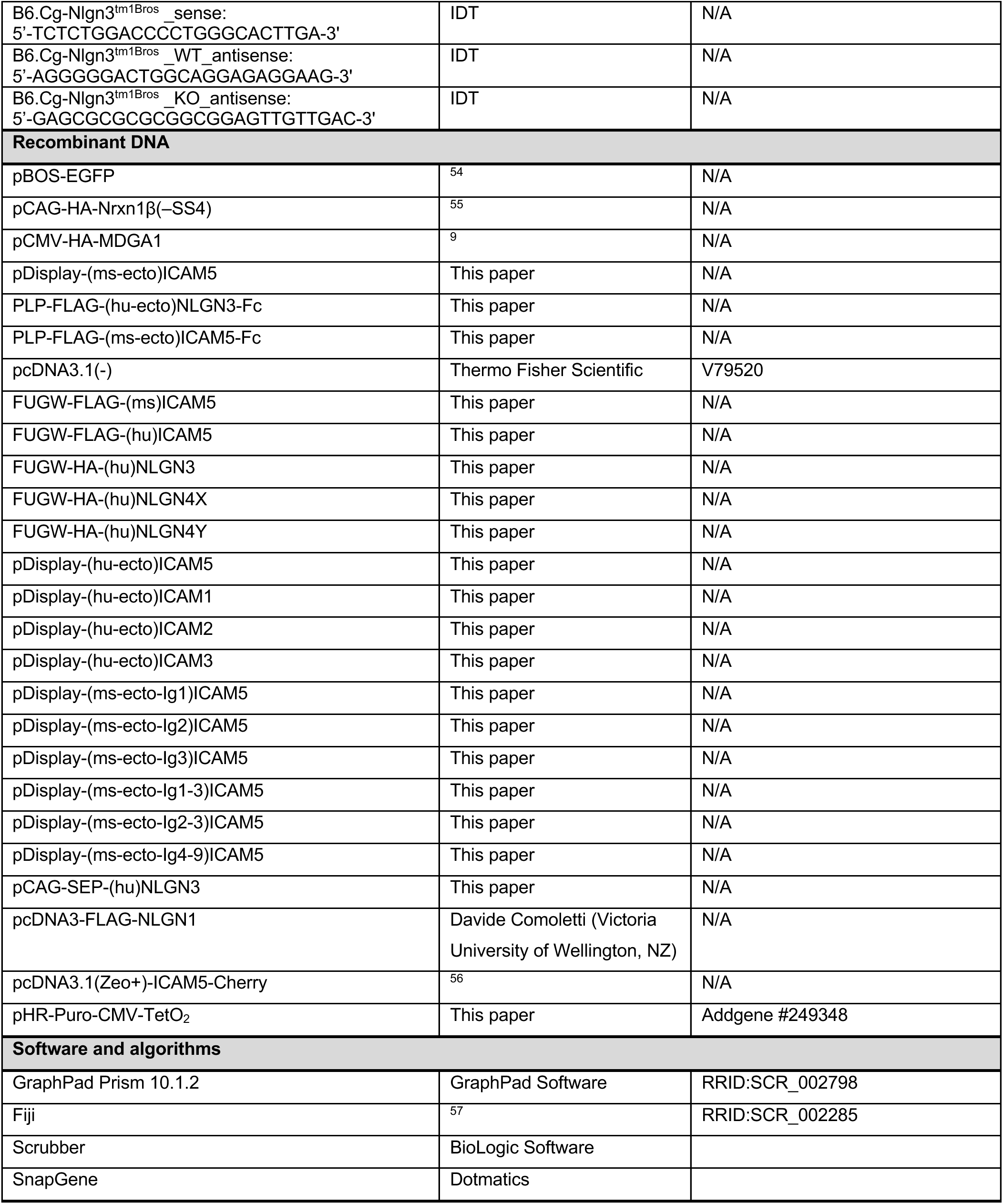

## EXPERIMENTAL MODELS AND SUBJECT DETAILS

### Animals

All animal experiments were conducted in accordance with local and national ethical guidelines and were approved by the respective Ethical Committees. Mice and rats were maintained under standard housing conditions with continuous access to food and water. The health and welfare of the animals were supervised by a designated veterinarian. The animal facilities comply with all appropriate standards (cage type, space per animal, temperature, light, humidity, food, and water), and all cages were enriched with materials that allow the animals to express their natural behaviors. Animals were maintained on a diurnal 12-hour light/dark cycle. For neuronal culture preparation, embryos were decapitated, and adult females were sacrificed by an overdose of isoflurane followed by cervical dislocation (mice) or decapitation (rats). Both male and female embryos were used to prepare the cultures. To the best of our knowledge, we are not aware of any influence of sex on the parameters analyzed in this study. The following mouse lines were used in this study: B6.Cg-Nlgn3^tm1Bros^ (*Nlgn3* full KO) and C57BL/6J^Icam5(-/-)^ (*Icam5* full KO). Wistar Han IGS rats were also used for the preparation of rat neuronal cultures and synaptosomes.

### Neuronal Cultures

Cortical neurons were cultured from E18–E19 Wistar rat embryos, B6.Cg-Nlgn3^tm1Bros^ mouse embryos, or C57BL/6J^Icam5(-/-)^ mouse embryos, as previously described ^58^. In the case of *Nlgn3* or *Icam5*, we used heterozygous females that had been crossed with the corresponding heterozygous males. After sacrificing the heterozygous females, embryos were removed, tissue was collected for genotyping, and the dissected cortices were maintained in Hibernate-E Medium (Thermo Fisher Scientific, cat. no. A1247601) supplemented with B27 (1:50; Thermo Fisher Scientific, cat. no. 17504044) during genotyping. Genotyping was performed using the KAPA HotStart Mouse Genotyping Kit (Roche, cat. no. KK7352). Following genotyping, cortical neurons were obtained from *Nlgn3* or *Icam5* KO embryos and the corresponding WT littermates. In brief, after cell dissociation, neurons were plated on coated glass coverslips (Glaswarenfabrik Karl Hecht, cat. no. 41001118) in 60-mm culture dishes (final density: 4×10^5^–7×10^5^ cells per dish), containing MEM (Thermo Fisher Scientific, cat. no. 11095080) supplemented with 10% (vol/vol) horse serum (Thermo Fisher Scientific, cat. no. 26050088) and 0.6% (wt/vol) glucose. Once neurons attached to the substrate (after 2–4 h), the coverslips were flipped over an astroglial feeder layer in 60-mm culture dishes containing neuronal culture medium: Neurobasal medium (Thermo Fisher Scientific, cat. no. 21103049) supplemented with B27 (1:50; Thermo Fisher Scientific, cat. no. 17504044), 12 mM glucose, GlutaMAX (1:400; Thermo Fisher Scientific, cat. no. 35050061), penicillin/streptomycin (1:500; Thermo Fisher Scientific, cat. no. 15140122), 25 μM β-mercaptoethanol, and 20 μg/mL insulin (Sigma-Aldrich, cat. no. I9278). Neurons grew face-down over the feeder layer but were kept separate from the glia by wax dots on the neuronal side of the coverslips. To prevent overgrowth of glia, neuron cultures were treated with 10 μM 5-fluoro-2′-deoxyuridine (Sigma-Aldrich, cat. no. F0503) after 3 days. Cultures were maintained in a humidified incubator with 5% (vol/vol) CO_2_/95% (vol/vol) air at 37 °C, feeding the cells once per week by replacing one-third of the medium in the dish.

### Cell lines

HEK293T-17 human embryonic kidney cells were obtained from the American Type Culture Collection (ATCC, cat. no. CRL-11268). HEK293T-17 cells were grown in DMEM (Thermo Fisher Scientific, cat. no. 11965092) supplemented with 10% (vol/vol) FBS (Thermo Fisher Scientific, cat. no. 10270106) and 1% (vol/vol) penicillin/streptomycin (Thermo Fisher Scientific, cat. no. 15140122).

## METHOD DETAILS

### Expression constructs and molecular cloning

The following list details the constructs used in this work, and their corresponding domain boundaries. Human intercellular adhesion molecule 1 (ICAM1; UniProt access code: P05362; Boundaries: Q28-E480), human intercellular adhesion molecule 2 (ICAM2; P13598; K25-Q223), human intercellular adhesion molecule 3 (ICAM3; P32942; Q30-H485), human intercellular adhesion molecule 5 (ICAM5; Q9UMF0; E32-P834), human Neuroligin-1 (NLGN1; Q8N2Q7; Q46-D635), human Neuroligin-2 (NLGN2; Q8NFZ4; E38-H612), human Neuroligin-3 (NLGN3; Q9NZ94; A38-D636), human Neuroligin-4X (NLGN4 X-linked; Q8N0W4; Q42-E602), human Neuroligin-4Y (NLGN4 Y-linked or NLGN5; Q8NFZ3; Q42-E602), human Neurexin-1 beta (β-NRXN1-LNS6(–SS4); P58400; G85-V295), human MAM domain-containing glycosylphosphatidylinositol anchor protein 1 (MDGA1; Q8NFP4; Q19-K925), and human Receptor-type tyrosine-protein phosphatase delta (PTPRD-Ig123-Fn1(+meA3-meB; E21-Q415).

Expression constructs were generated using the NEBuilder HiFi DNA Assembly Master Mix Kit (New England Biolabs, cat. no. E2621L) by inserting PCR-derived DNA fragments or gBlocks into the final vectors digested with the appropriate restriction enzymes. For FUGW-FLAG-(ms)ICAM5, FUGW-FLAG-(hu)ICAM5, FUGW-HA-(hu)NLGN3, FUGW-HA-(hu)NLGN4X, FUGW-HA-(hu)NLGN4Y, PCR fragments or gBlocks containing the full-length WT sequences were inserted into the pFUGW vector digested with AgeI and EcoRI. HA or FLAG epitope tags were incorporated immediately after the signal peptides. For PLP-FLAG-(ms-ecto)ICAM5-Fc and PLP-FLAG-(hu-ecto)NLGN3-Fc, PCR fragments or gBlocks containing the WT extracellular portions were inserted into the PLP vector digested with NotI and SaII. Finally, for pDisplay-(ms-ecto)ICAM5, pDisplay-(hu-ecto)ICAM5, pDisplay-(hu-ecto)ICAM1, pDisplay-(hu-ecto)ICAM2, pDisplay-(hu-ecto)ICAM3, pDisplay-(ms-ecto-Ig1)ICAM5, pDisplay-(ms-ecto-Ig2)ICAM5, pDisplay-(ms-ecto-Ig3)ICAM5, pDisplay-(ms-ecto-Ig1–3)ICAM5, pDisplay-(ms-ecto-Ig2–3)ICAM5, and pDisplay-(ms-ecto-Ig4–9)ICAM5, PCR fragments containing the full-length or partial extracellular domains were inserted into BglII- and SacII-digested pDisplay in frame with the signal peptide and HA epitope tag, and positioned upstream of the PDGFR transmembrane domain, allowing membrane anchoring of the extracellular portions of the expressed proteins.

All DNA constructs used in this study were sequence-verified. A complete list of plasmids is provided in the KEY RESOURCES TABLE.

### Expression and purification of recombinant proteins for cell surface binding assays and ECD-Fc affinity purifications

NLGN3_ECD_-Fc and ICAM5_ECD_-Fc proteins were produced by transient transfection of HEK293T cells using PEI (Polysciences, cat. no. 24765). 6 h after transfection, the medium was replaced by Opti-MEM (Thermo Fisher Scientific, cat. no. 22600134), and conditioned media were harvested 5 d later. Conditioned media were centrifuged, sterile-filtered, and applied to a CaptivA HF Protein A Affinity Resin (Repligen, cat. no. CA-HF) fast-flow column. After washing with wash buffer (50 mM HEPES (pH 7.4), 300 mM NaCl), bound proteins were eluted with IgG elution buffer (Thermo Fisher Scientific, cat. no. 21004). Fractions containing the proteins were pooled and dialyzed into PBS (137 mM NaCl, 2.7 mM KCl, 1.8 mM KH_2_PO_4_, and 10 mM Na_2_HPO_4_; pH 7.4) using a Slide-A-Lyzer (Thermo Fisher Scientific, cat. no. 66807), then concentrated using Amicon Ultra centrifugal units (Millipore, cat. no. UFC90500). The integrity and purity of the purified ecto-Fc proteins were assessed by SDS-PAGE followed by Coomassie staining, and protein concentrations were determined using a Bradford assay (Bio-Rad, cat. no. 5000202).

### Fc-ectodomain affinity capture of binding partners from synaptosome extracts

Affinity purification of extracellular binding partners was performed essentially as described previously for Ecto-Fc interactome screens ^27^. Briefly, whole brains from postnatal day 21–23 rats were homogenized on ice in extraction buffer supplemented with protease inhibitors using a Dounce homogenizer. Detergent extracts were prepared by addition of Triton X-100 (final concentration 1%) and incubated at 4 °C for ≥2 h with gentle stirring to solubilize membrane proteins. Insoluble material was removed by ultracentrifugation at 100,000 × g for 1 h at 4 °C, and the clarified detergent extract was used for affinity purification. For each purification, 12–13 mL of clarified extract was incubated overnight at 4 °C with 50–100 µg of Fc-fused ectodomain bait protein immobilized on Protein A affinity resin with end-over-end rotation. Equivalent purifications using Fc alone were performed in parallel as negative controls. Following incubation, bead suspensions were transferred to chromatography columns and extensively washed with high-salt wash buffer to remove nonspecific interactors, followed by a final low-salt wash. Bound proteins were eluted using acidic elution buffer and collected in microcentrifuge tubes. Eluted proteins were precipitated by addition of trichloroacetic acid (TCA) to a final concentration of 20% and incubated on ice overnight. Protein pellets were recovered by centrifugation, washed twice with ice-cold acetone, air-dried, and subsequently processed for tryptic digestion and MS analysis.

### Mass spectrometry (MS)

MS analysis of the NLGN3_ECD_-Fc and ICAM5_ECD_-Fc affinity-purified material was performed as previously described ^27,59^. For Orbitrap Fusion Tribrid MS analysis, tryptic peptides were purified using Pierce C18 spin columns (Thermo Fisher Scientific). Two to three micrograms of peptide were loaded via autosampler using a Thermo EASY-nLC 1200 UPLC onto a vented Acclaim PepMap 100 trap column (75 μm × 2 cm, nanoViper), followed by separation on a nanoViper analytical column (Thermo Fisher Scientific, 3 μm, 100 Å, C18, 75 μm × 500 mm) equipped with a stainless-steel emitter on the Nanospray Flex Ion Source at a spray voltage of 2.0 kV. Buffer A consisted of 94.875% (vol/vol) H_2_O, 5% (vol/vol) acetonitrile, and 0.125% (vol/vol) formic acid; buffer B consisted of 99.875% (vol/vol) acetonitrile and 0.125% (vol/vol) formic acid. The chromatographic separation was 4 h in total, with the following gradient: 0–7% B over 7 min, 7–10% over 6 min, 10–25% over 160 min, 25–33% over 40 min, 33–50% over 7 min, 50–95% over 5 min, and held at 95% for 15 min. Additional MS parameters were as follows: ion transfer tube temperature, 300 °C; Easy-IC internal mass calibration enabled; default charge state, 2; cycle time, 3 s. Full MS scans were acquired in the Orbitrap at a resolution of 60,000 with wide quadrupole isolation. The scan range was 300–1,500 m/z, maximum injection time 50 ms, AGC target 2 × 10^5^, one microscan, and S-lens RF level 60. Source fragmentation was disabled, and data were collected in positive centroid mode. Monoisotopic precursor selection was enabled for ions of charge +2 to +6 (unassigned charge states were rejected). Dynamic exclusion was enabled and set to 30 s (5 s exclusion duration) at ±10 ppm. Precursors were selected using the top-20 most intense ions, with an isolation window of 1.6 m/z, normal scan range, first mass 110 m/z, and collision energy 30%. For CID, MS/MS spectra were acquired in the ion trap with a scan rate of “rapid”, resolution equivalent to 30k, maximum injection time 75 ms, AGC target 1 × 10^4^, and Q set to 0.25. Ions were injected for all available parallelizable time.

Spectrum raw files were converted to ms1 and ms2 formats using RawXtractor or RawConverter (https://www.scripps.edu/yates/Software.html). Tandem mass spectra were searched against the UniProt rat proteome (downloaded 04-01-2013) using ProLuCID (version 3.1) on an Intel Xeon Linux cluster. The search space included all fully and half-tryptic peptides within the mass tolerance window, with no miscleavage constraints. Carbamidomethylation of cysteine (+57.02146 Da) was set as a static modification. Peptide-spectrum match (PSM) validity was assessed using DTASelect2, using SEQUEST-defined parameters XCorr and DeltaCN. Search results were grouped by charge state (+1, +2, +3, and >+3) and by tryptic status (fully, half-, and nontryptic), producing 12 subgroups. For each subgroup, distributions of XCorr, DeltaCN, and DeltaMass for direct and decoy PSMs were obtained and separated by discriminant analysis. Because complete separation is not possible, peptide match probabilities were calculated using a nonparametric fit of the score distributions. A minimum peptide confidence threshold of 0.95 was applied. The false discovery rate (FDR) was calculated as the proportion of reverse-decoy PSMs among all accepted PSMs. Proteins required at least one half-tryptic peptide of excellent quality (FDR < 0.001). For high-resolution data, a DeltaMass ≤ 10 ppm was also required. After filtering, protein-level FDRs were <1% for all samples.

### Expression and purification of recombinant proteins for SPR

Constructs were generated via seamless In-Fusion assembly cloning (Takara Bio) and were verified with either Sanger sequencing or whole-plasmid sequencing (MWG Eurofins, Source Genomics).

To make polyclonal stable HEK293 expression cell lines, cDNAs were cloned into a modified pHR-CMV-TetO_2_ lentiviral plasmid ^60^. In the latter, a puromycin antibiotic selection marker cDNA under control of a human elongation factor-1 alpha core (EF-1α_CORE_) promoter was introduced (yielding pHR-Puro-CMV-TetO_2_), allowing antibiotic selection of cells independently of the expression of the gene of interest (GOI). The cDNAs were fused C-terminally either with mVenus-Twinstrep-Avi-His6 tags for all ICAM5 constructs or with Avi-His6 tags for all other constructs.

Polyclonal stable expression cell lines were generated via lentiviral transduction of HEK293T or HEK293T TetR_LENTI_ cells using established protocols ^60,61^. Lentiviruses were generated by transfecting HEK293T Lenti-X packaging cells (Takara Bio) with a 1:1:1 w/w/w DNA plasmid mix consisting of pMD2.G (envelope plasmid), psPAX2 (packaging plasmid) and the pHR-Puro-CMV-TetO_2_ transfer plasmid encoding the GOI. After 72 h, virus-containing supernatants were harvested and filtered on a 0.45 μm PES filter. Polybrene (Merck) was added to a final concentration of 10 μg/mL to increase transduction efficiency, and expression cells were transduced. Transduced expression cells were first selected with the antibiotic puromycin (10 μg/mL) for up to 5–7 d, starting from 48 h post-infection. Selective media and antibiotic-free media were alternated and refreshed throughout the selection procedure, and cell death and expansion of resistant cell populations was monitored daily. For large-scale secreted protein production, confluent polyclonal stable cell lines were expanded into adherent roller bottle format (Greiner Bio-One). In the case of HEK293T TetR_LENTI_ cells, doxycycline (Dox) was added to 1 μg/mL to induce expression at the desired time point. Proteins were expressed for 5 d.

Conditioned expression media were collected and centrifuged at 4,000 g for 20 min at 20 °C, followed by sterile filtration on a 0.22 μm bottle-top filter. Filtered media were then concentrated and buffer exchanged to 100 mM Tris pH 8.0, 500 mM NaCl (TBS) on an ÄKTA Flux 6 tangential flow filtration system (Cytiva) and supplemented with 20 mM imidazole. Protease inhibitor cocktail tablets (cOmplete, EDTA-free, Roche Cat. No. 11873580001) were added to prevent protein degradation.

Proteins were purified by immobilized metal-affinity chromatography (IMAC) on an ÄKTA Pure 25M system (Cytiva) by loading the concentrated and buffer exchanged media onto a 5 mL HisTrap HP nickel column (Cytiva) pre-equilibrated with TBS buffer. The column was washed with TBS containing 50 mM imidazole, and protein was eluted with TBS containing 250 mM imidazole. Eluted proteins were immediately desalted into 20 mM HEPES pH 7.4, 150 mM NaCl (HBS) buffer.

3C-cleavable purification tags were removed during overnight digestion with recombinant human rhinovirus (HRV) 3C protease (1:500 protease:protein w/w ratio) at 20 °C, after which the protein mixture was reapplied to a 1 mL HisTrap HP nickel column (Cytiva) to remove His-tagged 3C protease, cleaved tags, uncleaved protein and contaminants. The HisTrap HP flowthrough containing pure, cleaved protein was concentrated and further purified by size-exclusion chromatography (SEC) on a Superdex 200PG HiLoad 16/600 PG column (Cytiva) equilibrated with HBS. Purified proteins were concentrated, snap-frozen in liquid nitrogen and stored at −80 °C. Protein quality (stability, aggregation, degradation) was monitored using fluorescence-based thermal denaturation (NanoTemper Tycho).

### Surface Plasmon Resonance (SPR)

For small-scale transient expression in HEK293 cells, cDNAs were cloned into the pHLsec-Avitag3 plasmid ^62^, resulting in proteins carrying a C-terminal biotin ligase (BirA) recognition site (Avitag). For small-scale production of *in vivo* biotinylated Avi-tagged proteins, HEK293T cells stably expressing ER-resident BirA (BirA-ER) ^60^ were transiently transfected. After 72 h of expression in the presence of 100 μM D-biotin, conditioned media were collected and dialyzed against 10 mM Tris (pH 7.4), 150 mM sodium chloride, 3 mM calcium chloride and 0.005% (vol/vol) Tween-20 (TBS-CT).

SPR experiments were performed on a Biacore T200 machine (GE Healthcare) operated at 25 °C and at a data collection frequency of 10 Hz; i.e. a temporal resolution of 0.1 s. Streptavidin (Sigma-Aldrich; prepared in 10mM sodium acetate (pH 4.5)) was chemically coupled via amine coupling chemistry onto CM5 chips (Cytiva) to a response unit (RU) level of 5000 RU. Then, the biotinylated proteins (ligands) were captured to the desired RU level (according to their respective molecular weight and desired binding responses). Every two analyte binding cycles, a buffer injection was performed, allowing for double referencing of the binding responses. SPR running buffer composition was 10 mM Tris pH 7.5, 150 mM sodium chloride, 0.005% (vol/vol) Tween-20, 3 mM calcium chloride and 1g/L bovine serum albumin (BSA; yielding TBS-CTB buffer). A serial dilution was prepared in the above buffer for each analyte by two-fold dilution and the subsequent solutions were passed over the chip at a flow rate of 25 μL/min. Each analyte was injected for 150 s, followed by a 240 s dissociation phase. For MDGA1, a 35 s regeneration step was included after the dissociation phase. The composition of regeneration solution used was 1 M magnesium chloride. Due to the high protein consumption associated with these assays, data are from an n = 1 experiment.

β-NRXN1(–SS4)_LNS6_ (500 RU), ICAM5_ECD_ (2200 RU), MDGA1_ECD_ (2000 RU) and PTPRδ(+meA3–meB)_Ig123-Fn1_ (860RU) were immobilized and NLGN3_ECD_ was injected as a twofold dilution series (100 μM to 12.2 nM, 14 concentrations total).

NLGN1_ECD_, NLGN2_ECD_, NLGN3_ECD_, NLGN4X_ECD_ and NLGN4Y_ECD_ (1000 RU) were immobilized and ICAM5 _ECD_, MDGA1 _ECD_ and β-NRXN1(–SS4)_LNS6_ were injected as a twofold dilution series (50 μM to 12.2 nM, 13 concentrations total).

Fitting and analysis of equilibrium binding data (1:1 or 1:2 Langmuir binding models) was performed with the program Scrubber (v2, BioLogic Software).

### Cell surface binding assay

HEK293T cells (40,000–60,000 cells/well) were plated on poly-D-lysine–coated 8-well chamber slides (Thermo Fisher Scientific, cat. no. 177402PK) in DMEM (Thermo Fisher Scientific, cat. no. 11965092) supplemented with 10% (vol/vol) FBS (Thermo Fisher Scientific, cat. no. 10270106) and 1% (vol/vol) penicillin-streptomycin (Thermo Fisher Scientific, cat. no. 15140122). At ∼50% confluency, cells were transfected with plasmids encoding the indicated receptors or EGFP using FuGENE 6 (Promega, cat. no. E2691) or PEI (Polysciences, cat. no. 24765) according to the manufacturer’s instructions. Twenty-four hours after transfection, live cells were incubated for 1 h at room temperature with Fc or ecto-Fc fusion proteins (10 µg/mL) diluted in DMEM containing 20 mM HEPES (pH 7.4), washed twice with DMEM/20 mM HEPES (pH 7.4), and fixed in 4% (wt/vol) paraformaldehyde and 4% (wt/vol) sucrose in PBS (137 mM NaCl, 2.7 mM KCl, 1.8 mM KH_2_PO_4_, and 10 mM Na_2_HPO_4_; pH 7.4) for 15 min at room temperature. After three washes with PBS, cells were permeabilized for 5 min at 4 °C with 0.25% (vol/vol) Triton X-100 in PBS. Cells were then blocked for 1 h in 10% (wt/vol) BSA in PBS and incubated overnight at 4°C with primary antibodies against the respective epitope tags diluted in 3% (wt/vol) BSA in PBS, followed by fluorophore-conjugated secondary antibodies (Jackson ImmunoResearch or Invitrogen) for 1 h at room temperature diluted in 3% (wt/vol) BSA in PBS.

### Co-culture assay

Co-culture assays were performed as previously described ^54^. Briefly, HEK293T cells were maintained in DMEM (Thermo Fisher Scientific, cat. no. 11965092) supplemented with 10% (vol/vol) FBS (Thermo Fisher Scientific, cat. no. 10270106) and 1% (vol/vol) penicillin–streptomycin (Thermo Fisher Scientific, cat. no. 15140122) and transfected using FuGENE 6 (Promega, cat. no. E2691) according to the manufacturer’s instructions. Twenty-four hours after transfection, HEK293T cells were mechanically dissociated and co-cultured with DIV 7–9 WT mouse cortical neurons for 16 h.

### Neuron transfection

Cortical neurons were transfected either at day *in vitro* (DIV) 3–7 or DIV 10–14. A total of 0.5 μg of each DNA plasmid was used per coverslip. Briefly, DNA plasmids were diluted in Tris-EDTA buffer (10 mM Tris-HCl and 2.5 mM EDTA; pH 7.3), followed by the dropwise addition of CaCl_2_ solution (2.5 M CaCl_2_ in 10 mM HEPES; pH 7.2) to the plasmid DNA solution to give a final concentration of 250 mM CaCl_2_. This solution was subsequently added to an equal volume of HEPES-buffered solution (274 mM NaCl, 10 mM KCl, 1.4 mM Na_2_HPO_4_, 42 mM HEPES; pH 7.2) and vortexed gently for 3 s. The resulting mixture, containing precipitated DNA, was then added dropwise to the coverslips in a 12-well plate containing 250 μL of conditioned neuronal culture medium with kynurenic acid (2 mM), followed by incubation for 2 h in a 37 °C, 5% (vol/vol) CO_2_/95% (vol/vol) air incubator. After 2 h, the transfection solution was removed, and 1 mL of conditioned neuronal culture medium containing kynurenic acid (2 mM), slightly acidified with HCl (approximately 5 mM final concentration), was added to each coverslip. The plate was then returned to a 37 °C, 5% (vol/vol) CO_2_/95% (vol/vol) air incubator for 20 min. Finally, coverslips were transferred back to the original dish containing conditioned culture medium and maintained at 37 °C in 5% (vol/vol) CO_2_/95% (vol/vol) air for 4 d to allow expression of the constructs.

### Immunocytochemistry

Neurons were fixed for 10–15 min at room temperature in 4% (wt/vol) paraformaldehyde and 4% (wt/vol) sucrose in PBS (137 mM NaCl, 2.7 mM KCl, 1.8 mM KH_2_PO_4_, and 10 mM Na_2_HPO_4_; pH 7.4) and permeabilized for 5 min at 4 °C with PBS containing 0.25% (vol/vol) Triton X-100. Cells were then blocked for 1 h at room temperature in 10% (wt/vol) BSA in PBS and incubated overnight at 4 °C with primary antibodies diluted in 3% (wt/vol) BSA in PBS. After three washes in PBS, cells were incubated for 1 h at room temperature with fluorophore-conjugated secondary antibodies (Jackson ImmunoResearch or Invitrogen) diluted in 3% (wt/vol) BSA in PBS. Coverslips were mounted using ProLong Gold Antifade mounting medium (Thermo Fisher Scientific, cat. no. P36930). For live staining, live rat cortical neurons were incubated for 15 min at room temperature with rabbit anti-HA antibody (1:100; Cell Signaling Technology, cat. no. 3724) diluted in conditioned neuronal culture medium. Neurons were then fixed for 10 min at room temperature in 4% (wt/vol) paraformaldehyde and 4% (wt/vol) sucrose in PBS, washed several times with PBS, and blocked for 1 h at room temperature in 10% (wt/vol) BSA in PBS. Cells were subsequently incubated for 1 h at room temperature with anti-rabbit secondary antibody diluted in 3% (wt/vol) BSA in PBS. Following permeabilization, neurons were processed for immunocytochemistry as described above.

### Preparation of neuronal lysates

DIV 7 cortical neurons were washed twice with ice-cold PBS (137 mM NaCl, 2.7 mM KCl, 1.8 mM KH_2_PO_4_, and 10 mM Na_2_HPO_4_; pH 7.4). After the final wash, PBS was aspirated and cells were lysed on ice in lysis buffer (25 mM Tris-HCl pH 7.5, 125 mM NaCl, 5 mM EDTA, 1% (vol/vol) Triton X-100, 1% (wt/vol) sodium deoxycholate, protease inhibitor cocktail [Roche, cat. no. 11697498001], and phosphatase inhibitor cocktail [Roche, cat. no. 4906845001]). A total of 100 μL lysis buffer was added per well, and cells were scraped using a plastic cell scraper and transferred to microcentrifuge tubes. Lysates were incubated for 30 min at 4 °C, followed by centrifugation at 15,000 rpm for 20 min at 4 °C. The resulting supernatant was collected into new tubes, and the insoluble pellet was discarded. Protein concentration was determined using the BCA Protein Assay (Thermo Fisher Scientific, cat. no. 23227). Protein lysates were subsequently used for Western blotting. For the SUnSET assay, DIV 7 cortical neurons were incubated in 2.5 mL of fresh neuronal medium containing puromycin (Sigma-Aldrich, cat. no. P8833; 2 μM) for 45 min, followed by neuronal lysate preparation as described above.

### Western blotting

Protein extracts were denatured by boiling at 95 °C for 5 min with 5× Laemmli sample buffer containing 62.5 mM Tris-HCl pH 6.8, 2% (vol/vol) SDS, 10% (vol/vol) glycerol, 5% (vol/vol) β-mercaptoethanol, and 0.01% (wt/vol) bromophenol blue. Samples were resolved on 10% (wt/vol) polyacrylamide gels (Bio-Rad, cat. no. 4561034) in Tris–glycine SDS running buffer (25 mM Tris base, 192 mM glycine, and 0.1% vol/vol SDS; pH 8.3). Protein transfer was performed using the Trans-Blot Turbo Mini Nitrocellulose Transfer Kit (Bio-Rad, cat. no. 1704270) according to the manufacturer’s instructions. Membranes were blocked for 1 h at room temperature in 5% (wt/vol) low-fat milk prepared in Tris-buffered saline (300 mM NaCl, 25 mM Tris-HCl (pH 7.5), containing 0.05% (vol/vol) Tween-20; TBS- T). Primary antibodies were diluted in 5% (wt/vol) low-fat milk/TBS-T and incubated with membranes overnight at 4 °C. After five washes in TBS-T at room temperature, membranes were incubated for 1 h at room temperature with horseradish peroxidase-conjugated secondary antibodies. Membranes were then washed four times in TBS-T followed by one wash in low-salt TBS-T (150 mM NaCl, 25 mM Tris-HCl (pH 7.5), 0.05% (vol/vol) Tween-20) and incubated with chemiluminescent ECL substrates (Thermo Fisher Scientific, cat. no. 34095, cat. no. 34579, or cat. no. 32132) for up to 5 min. Signals were detected using an Amersham ImageQuant 800 Western blot imaging system (Cytiva). Membranes were stripped using Restore PLUS Western Blot Stripping Buffer (Thermo Fisher Scientific, cat. no. 46430) according to the manufacturer’s instructions. Western blot images were quantified using Fiji.

## QUANTIFICATION AND STATISTICAL ANALYSIS

### Image analysis and quantification

All images from immunocytochemistry experiments in fixed cells were acquired using either a Leica SP8 laser-scanning confocal microscope (Leica Microsystems) or a Zeiss LSM 710 confocal microscope (Carl Zeiss Microscopy). The same confocal acquisition settings were applied to all images obtained within the same independent experiment. Acquisition parameters were adjusted to avoid pixel saturation. Z-stack images were converted to maximum-intensity projections prior to analysis. Quantitative image analysis was performed using Fiji.

Cell surface binding was quantified by measuring the Fc signal associated with HEK293T cells. Briefly, HEK293T cells were identified based on GFP or HA immunostaining, and cell area was defined from thresholded images. The Fc signal was also thresholded, and the area of Fc labeling overlapping each HEK293T cell was measured. Binding was expressed as the ratio of Fc-labeled area to the total area of the corresponding HEK293T cell.

For quantification of total dendritic length, custom macros were implemented in Fiji. Only dendrites, defined as GFP-positive neurites co-labeled with MAP2, were included in the analysis. GFP images were thresholded to generate binary masks of the dendritic arbor and processed with the 2D/3D Skeletonize function to obtain single-pixel-wide skeletons. Skeletonized images were converted to maximum-intensity projections, and total dendritic length was then measured using the Analyze Skeleton (2D/3D) function.

Fluorescence intensity of phospho-PAK and F-actin was quantified from thresholded images. Growth cone regions were defined by phalloidin staining (F-actin) within MAP2-positive neurites. For each cell, the integrated fluorescence intensity of phospho-PAK (or F-actin) within these regions was summed and normalized to the corresponding F-actin-positive area from the respective thresholded images.

Fluorescence intensity of PSD-95 was quantified from thresholded images. Representative dendritic regions were selected using GFP-positive neurites co-labeled with MAP2 as guides. For each cell, the sum of the integrated PSD-95 fluorescence intensity within these dendritic regions was measured and normalized to the corresponding dendritic area defined from thresholded GFP images.

Accumulation of Synapsin1, used as a proxy for synaptogenesis, was quantified by measuring the Synapsin1 signal associated with NLGN-expressing HEK293T cells. HEK293T cells were identified based on NLGN-positive immunostaining, and cell area was defined from thresholded images. Synapsin1 labeling was also thresholded, and the area of Synapsin1 signal overlapping each HEK293T cell was measured. Synapsin1 accumulation was expressed as the ratio of Synapsin1-positive area to the total area of the corresponding HEK293T cell.

### Statistics

Results are presented as boxplots (median, 25^th^–75^th^ interquartile range, and minimum to maximum) or as individual data points with mean ± SEM. Normality of data distributions was assessed using the Shapiro–Wilk test and statistical significance was evaluated using either a parametric unpaired t-test or nonparametric Mann–Whitney and Kruskal–Wallis (followed by Dunn’s multiple comparisons test) tests, as appropriate. Data from at least two independent experiments were included in each analysis. Statistical analyses were performed using GraphPad Prism version 10.1.2.

---

**Supplemental Figure 1.**
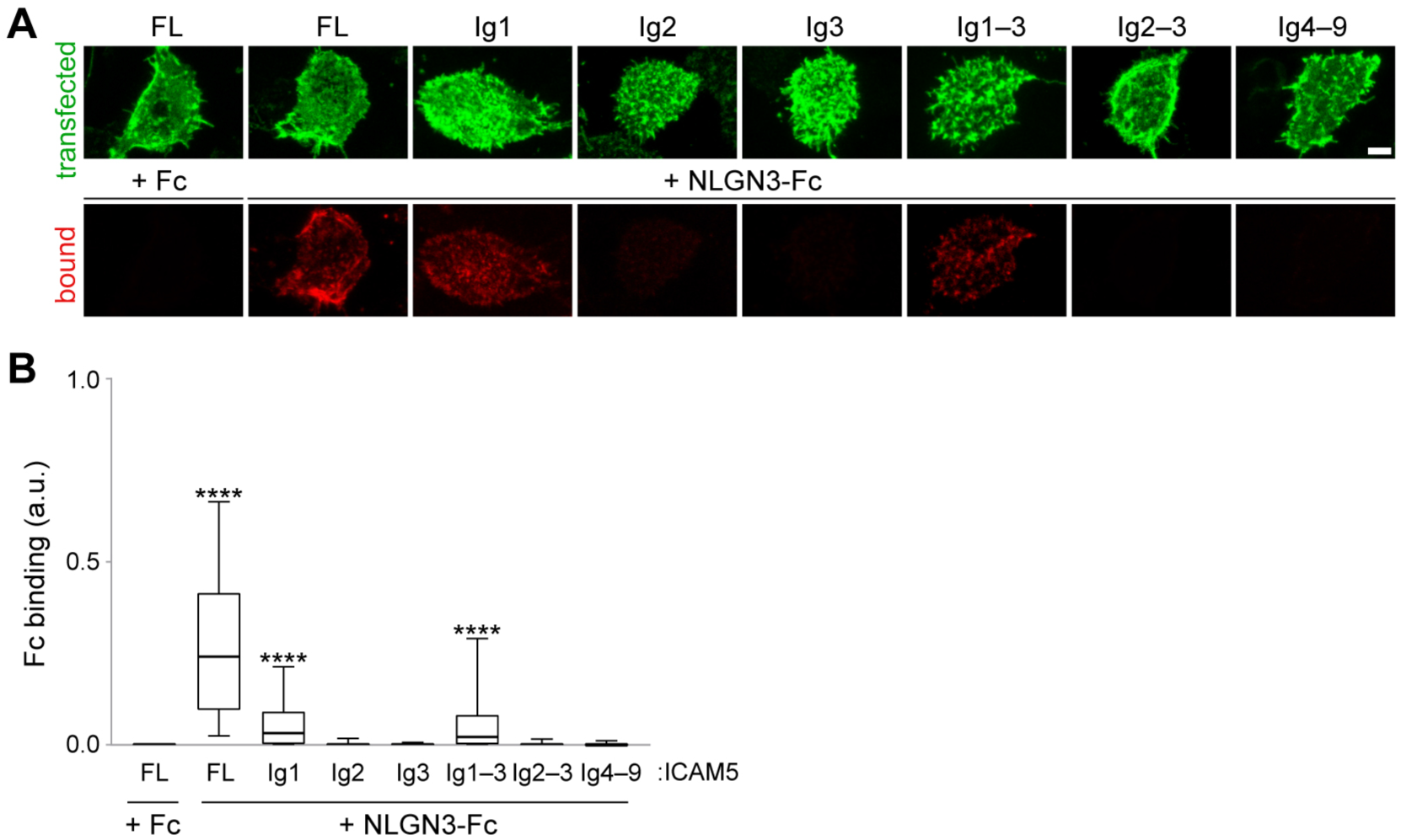
Ig domain 1 of ICAM5 is involved in NLGN3–ICAM5 interaction. (**A**) Cell-surface binding assays. NLGN3ECD-Fc (red) bound to HEK293T cells (green) expressing full-length (FL), Ig1 or Ig1–3 of HA-tagged ICAM5 ECD, but not to cells expressing Ig2, Ig3, Ig2–3, or Ig4–9. Scale bar, 5 µm. (**B**) Quantification of panel A: Fc clustering area normalized to the area of the respective HEK293T cell, as determined by HA staining (Fc binding). Data are shown as box plots (median, 25^th^–75^th^ percentile, and minimum to maximum). ****p < 0.0001 (Kruskal–Wallis test followed by Dunn’s multiple comparisons test, 3 independent experiments). a.u., arbitrary units.

**Supplemental Figure 2.**
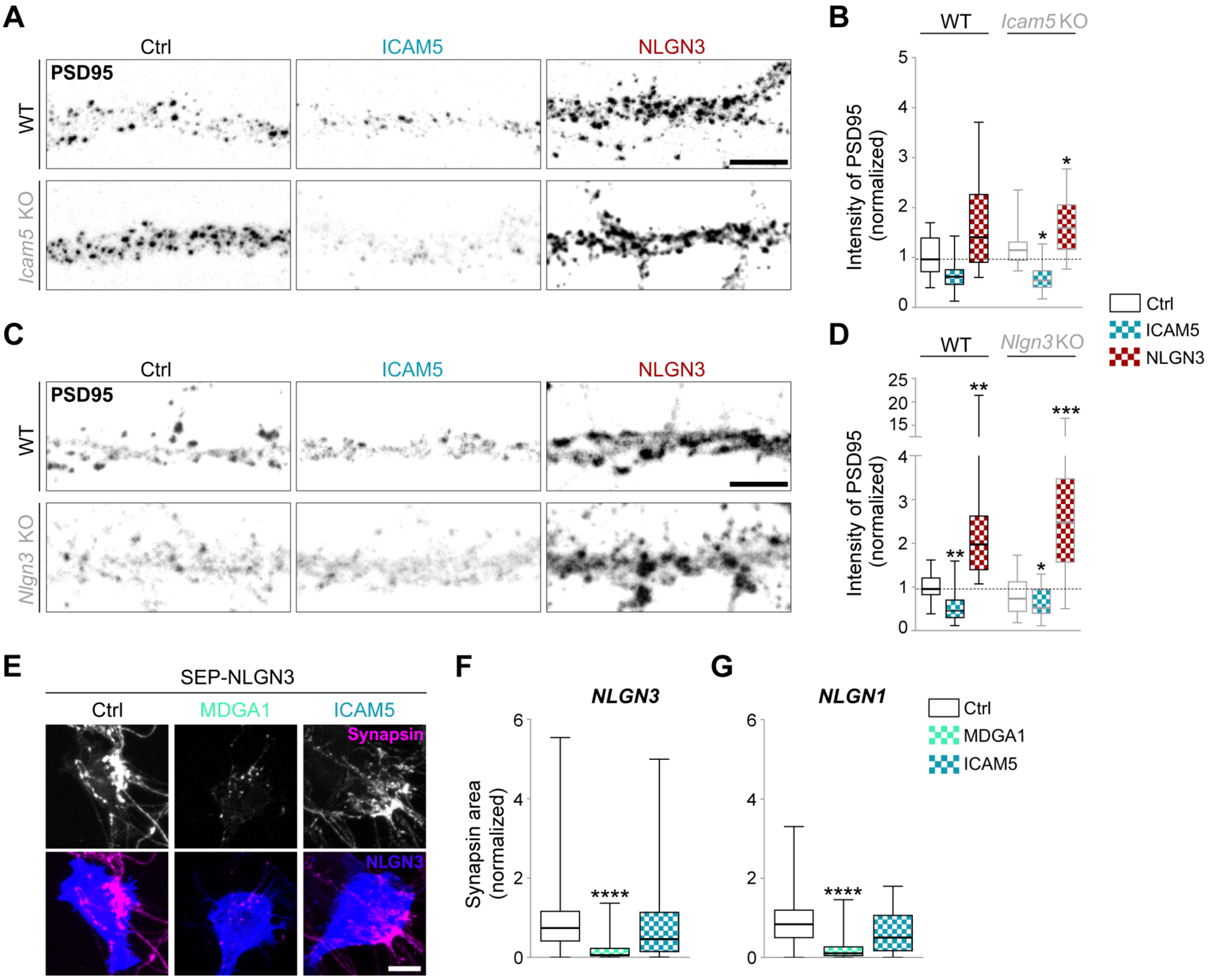
NLGN3–ICAM5 does not control synapse formation. (**A, C**) Representative images of DIV 14 WT or *Icam5* KO (A), or DIV 14 WT or *Nlgn3* KO (C) mouse cortical neurons transfected at DIV 10 with GFP and empty vector (Ctrl), GFP and FLAG-tagged ICAM5, or GFP and HA-tagged NLGN3. Neurons were immunostained for PSD-95 (grayscale), GFP (not shown), and HA or FLAG (not shown). Scale bar, 10 µm. (**B, D**) Quantification of panels A and C: total PSD-95 fluorescence intensity normalized to dendritic area, defined as GFP staining, and to control cells. Data are shown as box plots (median, 25^th^–75^th^ percentile, and minimum to maximum). *p < 0.05; **p < 0.01; ***p = 0.0005 (Kruskal–Wallis test followed by Dunn’s multiple comparisons test, 2 (panel B) or 3 (panel D) independent experiments). (**E**) HEK293T cells co-expressing Super Ecliptic pHluorin (SEP)-tagged NLGN3 with empty vector (Ctrl), HA-tagged MDGA1, or mCherry-tagged ICAM5 at a 1:5 ratio, were co-cultured with DIV 8–10 WT mouse cortical neurons. Co-cultures were immunostained for SEP (blue), Synapsin1 (grayscale and magenta), HA (not shown), and mCherry (not shown). Scale bar, 10 µm. (**F, G**) Quantification of Synapsin1 clustering area normalized to the area of the respective NLGN3-expressing (F) or NLGN1-expressing (G) HEK293T cell and to control cells. ****p < 0.0001 (Kruskal–Wallis test followed by Dunn’s multiple comparisons test, 3 independent experiments).

**Supplemental Figure 3.**
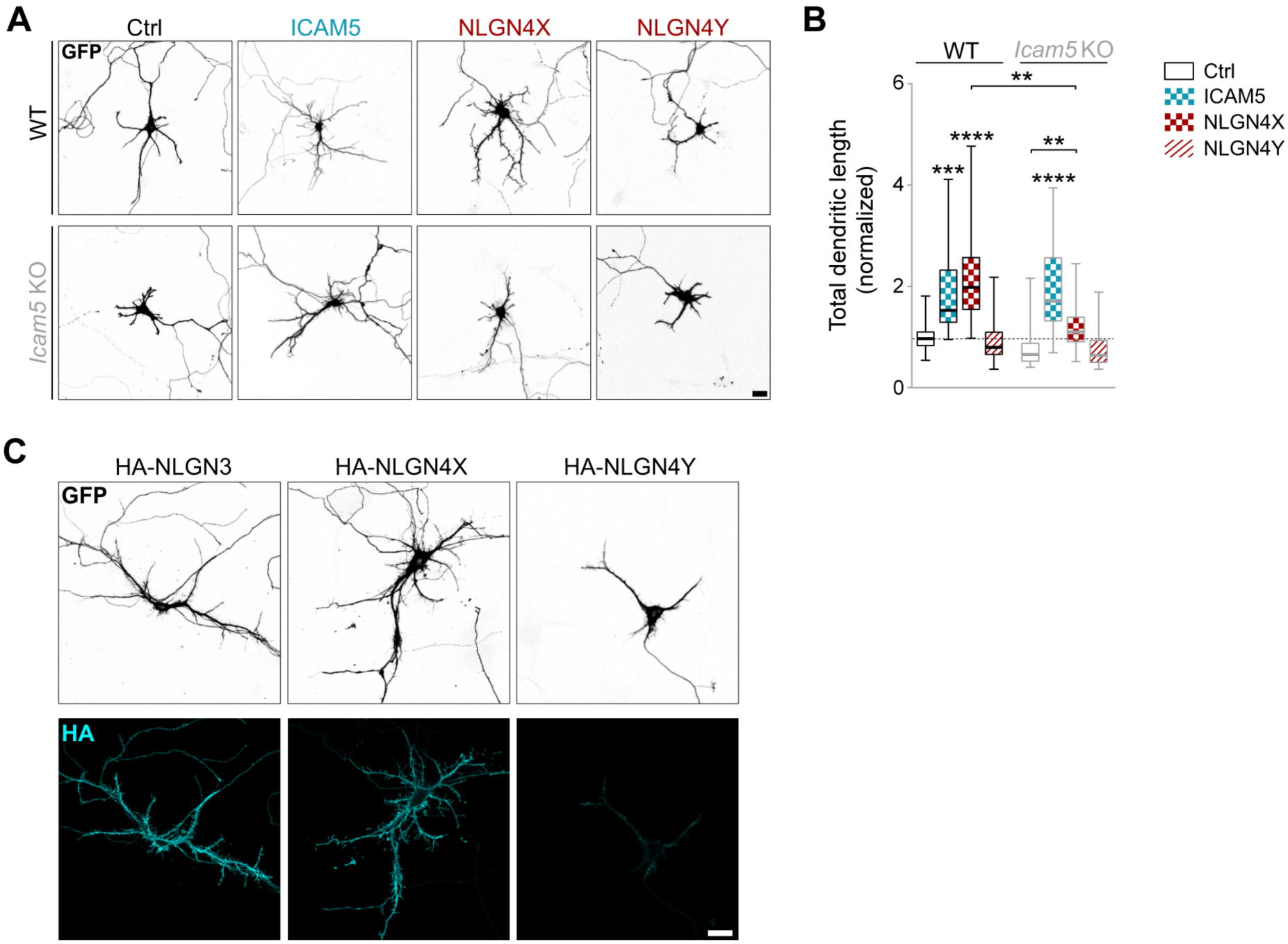
NLGN4X promotes dendritic outgrowth in an ICAM5-dependent manner. (**A**) Representative images of DIV 7 WT or *Icam5* KO mouse cortical neurons transfected at DIV 3 with GFP and empty vector (Ctrl), GFP and FLAG-tagged ICAM5, GFP and HA-tagged NLGN4X, or GFP and HA-tagged NLGN4Y. Neurons were immunostained for GFP (grayscale), MAP2 (not shown), and HA or FLAG (not shown). Scale bar, 20 µm. (**B**) Quantification of panel C: total dendritic length, defined as GFP signal in MAP2-positive neurites and normalized to control cells. Data are shown as box plots (median, 25^th^–75^th^ percentile, and minimum to maximum). **p < 0.01; ***p = 0.0001; ****p < 0.0001 (Kruskal–Wallis test followed by Dunn’s multiple comparisons test, 3 independent experiments). (**C**) Representative images of DIV 7 WT rat cortical neurons transfected at DIV 3 with GFP and WT HA-tagged NLGN3, NLGN4X or NLGN4Y. Neurons were immunostained for surface NLGN levels (cyan) and GFP (grayscale). Scale bar, 20 µm.

**Supplemental Figure 4.**
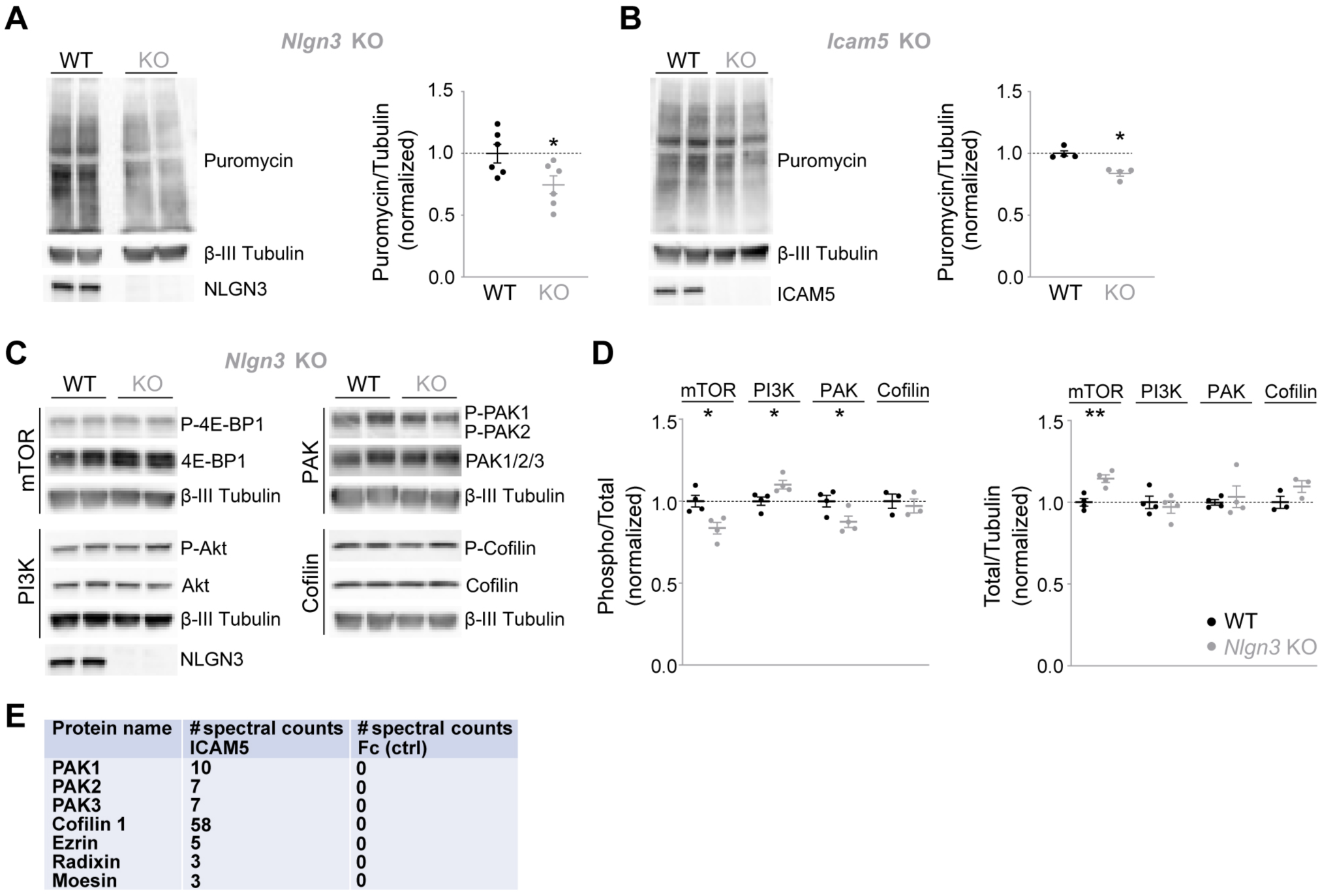
The NLGN3–ICAM5 complex regulates intracellular signaling pathways. (**A, B**) SUnSET assay in cortical neurons prepared from *Nlgn3* or *Icam5* KO and WT littermate embryos. DIV 7 WT or *Nlgn3* KO (A) and WT or *Icam5* KO (B) mouse cortical neurons were incubated with puromycin (2 µM, 45 min). Lysates were then prepared and analyzed by western blot using anti-puromycin, anti-βIII-tubulin (loading control), and anti-NLGN3 or anti-ICAM5 antibodies. Quantification of puromycin band intensities was normalized to βIII-tubulin and to WT control. Data are shown as individual data points with mean ± SEM. *p < 0.05 (unpaired t-test, 6 independent experiments; panel A) and *p < 0.05 (Mann–Whitney test, 4 independent experiments; panel B). (**C**) Western blot analysis of signaling pathways in DIV7 cortical neurons derived from WT or *Nlgn3* KO mice. Lysates were probed for markers of mTOR signaling (4E-BP1 and phospho-4E-BP1 [Thr37/46]), PI3K signaling (Akt and phospho-Akt [Ser473]), PAK signaling (total PAK1–3 and phospho-PAK1 [Ser144] / phospho-PAK2 [Ser141]), and the actin regulator Cofilin (total and phospho-Cofilin [Ser3]). βIII-tubulin was used as a loading control, and NLGN3 immunoblotting confirmed loss of NLGN3 expression in *Nlgn3* KO neurons. (**D**) Quantification of panel C: phospho-band intensities were normalized to the corresponding total protein levels and to WT control, while total protein band intensities were normalized to βIII-tubulin and to WT control. Data are shown as individual data points with mean ± SEM. *p < 0.05; **p < 0.01 (Unpaired t-test, 3-4 independent experiments). (**E**) PAK1–3, Cofilin 1, Ezrin, Radixin and Moesin proteins were identified with elevated spectral counts in the ICAM5ECD-Fc interactome performed using postnatal day 21-23 whole rat brain synaptosomes, and were not present in the negative control.

